# Inferring fine-scale mutation and recombination rate maps in aye-ayes (*Daubentonia madagascariensis*)

**DOI:** 10.1101/2024.12.28.630620

**Authors:** Vivak Soni, Cyril J. Versoza, John W. Terbot, Jeffrey D. Jensen, Susanne P. Pfeifer

**Author notes:** these authors contributed equally to this work (Vivak Soni, Cyril J. Versoza). co-corresponding authors; jointly supervised the project (;).

## Abstract

The rate of input of new genetic mutations, and the rate at which that variation is reshuffled, are key evolutionary processes shaping genomic diversity. Importantly, these rates vary not just across populations and species, but also across individual genomes. Despite previous studies having demonstrated that failing to account for rate heterogeneity across the genome can bias the inference of both selective and neutral population genetic processes, mutation and recombination rate maps have to date only been generated for a relatively small number of organisms. Here, we infer such fine-scale maps for the aye-aye (*Daubentonia madagascariensis*) – a highly endangered strepsirrhine that represents one of the earliest splits in the primate clade, and thus stands as an important outgroup to the more commonly-studied haplorrhines – utilizing a recently released fully-annotated genome combined with high-quality population sequencing data. We compare our indirectly inferred rates to previous pedigree-based estimates, finding further evidence of relatively low mutation and recombination rates in aye-ayes compared to other primates.

## Introduction

The rate of input of new genetic variation, and the rate at which that variation is shuffled into potentially novel combinations via crossover and non-crossover events, are fundamental evolutionary forces shaping observed genomic diversity. Over the past decades, it has become clear that mutation rates vary at a variety of scales, from between sites in a genome, to between individuals in a population, to between populations of a species, as well as broadly across the Tree of Life (see reviews of Baer et al. 2007; Lynch 2010; Hodgkinson and Eyre-Walker 2011; Pfeifer 2020b). The same is true of recombination, with modifications of underlying rates observed to occur at even more rapid timescales (see reviews of Ritz et al. 2017; Stapley et al. 2017). Importantly, heterogeneity in both mutation and recombination rates across a genome can significantly alter interactions between other evolutionary processes; for example, modifying Hill-Robertson effects (Hill and Robertson 1966; Felsenstein 1974), thereby modulating the genomic impact of selection at linked sites (Maynard Smith and Haigh 1974; Begun and Aquadro 1992; Charlesworth et al. 1993; and see Charlesworth and Jensen 2021, 2022). Furthermore, neglecting this underlying rate heterogeneity in favor of using single, species-averaged rates for mutation and recombination – as is common practice in evolutionary models – has been shown to result in potentially mis-leading inference when performing downstream analyses that rely on these estimates (e.g., for inferring both population history and distributions of fitness effects, Soni et al. 2024a; Soni and Jensen 2024; and see Dapper and Payseur 2018; Samuk and Noor 2022; Ghafoor et al. 2023).

Aside from classical disease-incidence approaches (e.g., Haldane 1932, 1935), there are generally two classes of experiments to infer mutation rates in primates and other large organisms. Direct mutation rate estimation relies on high-throughput genome sequencing of parent-offspring trios or multi-generation pedigrees, counting the number of *de novo* mutations occurring from one generation to the next (see review of Pfeifer 2020b). As mutations are rare, this generally results in only a genome-wide estimate over the limited number of generations considered, rather than providing a finer-scale map. Relatedly, tremendous caution must be exercised in the applied computational approach as errors introduced during sequencing will generally far outnumber genuine spontaneous mutations (Pfeifer 2021; Bergeron et al. 2022).

Alternatively, indirect mutation rate estimation from species-level divergence data instead relies on Kimura’s (1968) observation that the neutral mutation rate is equal to the neutral divergence rate. Specifically, the number of substitutions *K* that accumulate in a lineage in time *T* is equal to (μ/*G*)*T*, where μ is the per-generation mutation rate and *G* the generation time. As such, historically-averaged mutation rates can be inferred from phylogenetic sequence data in neutral genomic regions, with the caveat that such estimates must generally be couched within the context of underlying uncertainties in both divergence and generation times (thus generally resulting in a range of possible mutation rates). Complicating matters further, the identification of neutral regions necessary for this indirect rate estimation requires high-quality genome annotations which are not yet widely available for many organisms.

Similarly for recombination, taking a pedigree-based approach enables the detection of contemporary crossover and non-crossover events in males and females separately. As with direct mutation rate estimation, these approaches have the advantage of direct observation, though the genome-scale resolution is again relatively coarse given the small number of meiotic exchanges that can be observed within a pedigree (see the review of Clark et al. 2010). By contrast, population-based approaches using unrelated individuals can indirectly infer historical recombination rates from patterns of linkage disequilibrium (LD) observed in the sample (see reviews of Stumpf and McVean 2003; Peñalba and Wolf 2020). As such, these approaches offer a higher genomic resolution and may thus provide for fine-scale mapping, though inferred rates are necessarily sex-averaged, and may be confounded by other population-level factors that can alter levels of LD (e.g., population history or selective effects; Dapper and Payseur 2018; Samuk and Noor 2022). For this reason, it is important to both directly model a fit demographic history when performing such inference, and to carefully annotate neutral genomic regions prior to analysis (Johri et al. 2020, 2022).

In primates, many of the highest quality estimates of both mutation and recombination rates have been obtained in humans and their closest relatives (i.e., non-human great apes) as well as in species of biomedical relevance (e.g., Kong et al. 2002; Auton et al. 2012; Stevison et al. 2016; Pfeifer 2020a; Xue et al. 2020; Wall et al. 2022; Versoza, Weiss, et al. 2024). In humans, for example, large-scale sequencing of pedigrees has yielded mutation rate estimates of ∼10^−8^ per base pair per generation (see review of Ségurel et al. 2014), which is roughly two-fold lower than the initial indirect estimates obtained from phylogenetic data (Nachman and Crowell 2000; Kondrashov 2003); while crossover rates have been inferred to range from 0.96 cM/Mb to 2.11 cM/Mb for the longest and shortest autosomes, respectively, with an overall sex-averaged rate of ∼1 cM/Mb (Kong et al. 2002). Recently however, owing to the generation of high-quality population genomic data from pedigreed individuals, combined with the release of a fully annotated, chromosomal-level genome assembly (Versoza and Pfeifer 2024), we now additionally have direct mutation and recombination rate estimates for aye-ayes (*Daubentonia madagascariensis*), a highly-endangered strepsirrhine that represents one of the earliest splits in the primate clade (Versoza et al. 2024a,b; Versoza, Lloret-Villas, et al. 2024). These direct estimates suggested an average genome-wide mutation rate of ∼1.1 x 10^−8^ per base pair per generation for the species – although mutation rates in the wild may be closer to a rate of ∼0.4 x 10^−8^ per base pair per generation, as was estimated for individuals in the pedigree reproducing at an early age – and a sex-averaged crossover rate of 0.85 cM/Mb. Importantly, utilizing polymorphism data from unrelated individuals, Terbot et al. (2024) additionally estimated a well-fitting population history for aye-ayes (and see Soni et al. 2024b), describing a severe and relatively ancient population decline in the species coinciding with the arrival of humans to Madagascar, as well as a far more recent decline likely associated with habitat destruction and fragmentation over the past few decades.

Taking advantage of this newly available high-coverage genome-wide polymorphism data from both unrelated and pedigreed individuals, the recent annotation of the genome enabling the masking of functional (i.e., directly selected) regions, as well as these pedigree-based direct coarse-scale estimates allowing for meaningful comparison, we here infer indirect fine-scale mutation and recombination rate maps across the aye-aye genome utilizing both levels and patterns of variation as well as divergence from other closely related primate species. Aside from the biological insight into the rates of mutation and recombination gained in this study, by allowing for the incorporation of the observed rate heterogeneity, these newly developed fine-scale maps will thus also be vitally important to improve future primate evolutionary models.

## Results and Discussion

### Fine-scale mutation rate map

We calculated aye-aye divergence by removing the existing (but outdated) aye-aye genome from the 447-way multiple species alignment, consisting of the combined mammalian multiple species alignment of the Zoonomia Consortium (2020) and the primate multiple species alignment of Kuderna et al. (2024), and replaced it with the current NCBI reference genome for the species (i.e., the high-quality, fully annotated aye-aye genome of Versoza and Pfeifer (2024); see the “Materials and Methods” section for details). By masking both functional regions and segregating variants, we calculated neutral divergence across accessible sites for a range of window sizes (1kb, 10kb, 100kb, and 1Mb), yielding a mean neutral divergence rate of 0.043 at the 1Mb-scale relative to the reconstructed ancestor (Supplementary Figure S1). Utilizing lower– and upper-bounds of aye-aye divergence times (54.9 million years ago [mya] and 74.7 mya; Horvarth et al. 2008) and bounds of likely generation times (3 years and 5 years; Ross 2003; Louis et al. 2020), we calculated neutral mutation rates across these genomic windows, as depicted in Table 1. The average mutation rate varied from 1.73 x 10^−9^ mutations per base pair per generation (under a divergence time of 74.7 mya and a generation time of 3 years) and 3.93 x 10^−9^ mutations per base pair per generation (under a divergence time of 54.9 mya and a generation time of 5 years). Figure 1a provides density plots of mutation rates for these divergence and generation times, whilst Figures 1b and 1c provide the heterogeneity in mutation rates across a single chromosome-length scaffold (using the longest autosomal scaffold as an example; and see Supplementary Figures S2-S14 for mutation rate heterogeneity across all other autosomal scaffolds) and across the whole genome, respectively.

**Figure 1:**
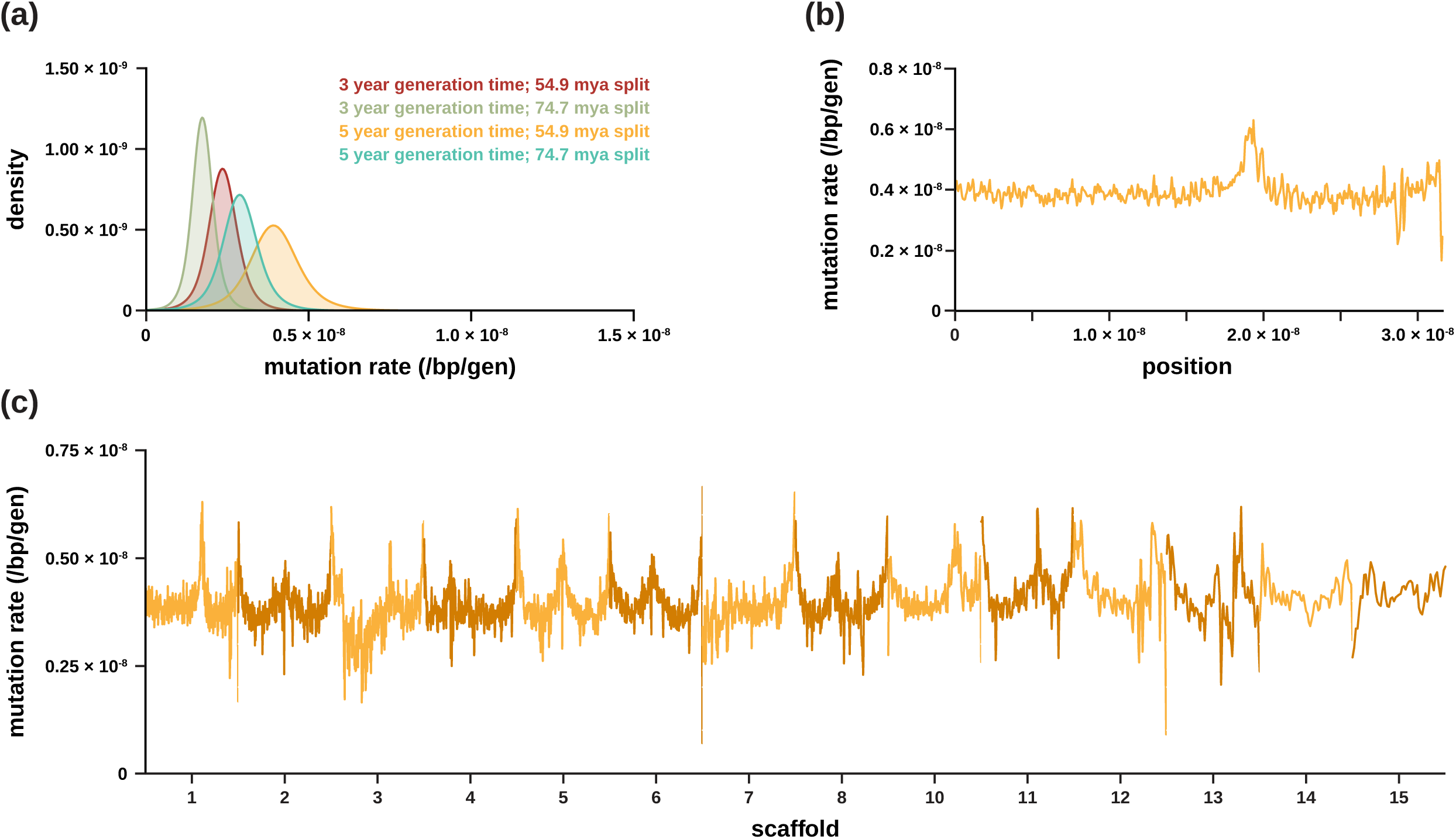
(a) Density plots of the per base pair per generation (/bp/gen) mutation rate implied by neutral divergence for two possible generation times (3 years and 5 years; Ross 2003; Louis et al. 2020) and two possible divergence times (54.9 million years ago [mya] and 74.7 mya; Horvarth et al. 2008). (b) Fine-scale mutation rates along the longest autosomal scaffold (i.e., scaffold 1) for genomic windows of size 1Mb, with a 500kb step size (see Supplementary Figures S2-14 for mutation rate heterogeneity across all other autosomal scaffolds). (c) Genome-wide mutation rates for genomic windows of size 1Mb, with a 500kb step size.

**Table 1:** Inferred aye-aye divergence times based on the observed mean neutral divergence rate of 0.043 for two different possible generation times (3 years and 5 years; Ross 2003; Louis et al. 2020) and three different pedigree-based mutation rates estimated for parents of differing ages by Versoza et al. (2024a) (shown in blue). Relatedly, the resulting divergence-based mutation rate estimates based on two possible divergence times (54.9 million years ago [mya] and 74.7 mya; Horvarth et al. 2008) and two possible generation times (3 years and 5 years; Ross 2003; Louis et al. 2020) are given for comparison (shown in orange).

|  |  | pedigree-based mutation rate |  |  | divergence time |  |
| --- | --- | --- | --- | --- | --- | --- |
|  |  | 4.0E-09 | 1.1E-08 | 2.0E-08 | 54.9 mya | 74.7 mya |
| generation | 3 | 32.3 mya | 11.7 mya | 6.45 mya | 2.36E-09 | 1.73E-09 |
| time (years) | 5 | 53.8 mya | 19.5 mya | 10.8 mya | 3.93E-09 | 2.89E-09 |

Taking the reverse tack, we additionally estimated aye-aye divergence times utilizing the recently inferred mutation rates from multi-generation aye-aye pedigree data (Table 1; Versoza et al. 2024a). These rates ranged from 0.4 x 10^−8^ per base pair per generation in individuals born to young parents (<12 years of age) to 2.0 x 10^−8^ per base pair per generation in individuals born to old parents (>24 years of age), with an average rate of ∼1.1 x 10^−8^ per base pair per generation, resulting in estimated divergence times spanning the very large range from 53.8 mya to 6.45 mya (when considering the highest and lowest generation times as well). These results strongly suggest that average ages of reproduction in the wild are comparatively young, given that the rates associated with older parents in captivity provide unrealistically recent divergence times relative to the fossil record (Gingerich 2006; Smith et al. 2006; and see the review of Gingerich 2012) – an observation in agreement with previous ecological studies that reported average reproductive ages of 3 to 5 years in the wild (Ross 2003; Louis et al. 2020). Further, the times associated with younger parents are consistent with previous estimates of divergence based on a limited set of genetic markers encompassing ∼9kb of nuclear sequence (Horvath et al. 2008), and thus the lower direct pedigree mutation rate of 0.4 x 10^−8^ per base pair per generation is likely the more appropriate long-term estimate for the species. Indeed, given that this estimate falls within our indirectly inferred mean mutation rate in this study as well, and that prosimians have been shown to have generally lower mutation rates compared to other primates (Tran and Pfeifer 2018; Chintalapati and Moorjani 2020), these results taken together represent a strong body of evidence that supports relatively low mutation rates in aye-ayes. Importantly, there is a considerable discordance in divergence time estimates of the strepsirrhine–haplorrhine split between those based on molecular data and the sparse fossil record – with the former placing the split as early as 90 mya and the latter at 55 mya (Hartwig 2011). Hence, with our improved estimates of mutation rates from both pedigree-based and divergence data, our estimate of ∼53.8 mya is in agreement with the origin of primates (Tavaré et al. 2002; Zhang et al. 2008), and thus with strepsirrhines representing one of the earliest splits in the primate clade (Pozzi et al. 2014).

### Fine-scale recombination rate map

We utilized two different approaches to infer fine-scale rates of recombination. The first, LDhat (McVean et al. 2002, 2004; Auton and McVean 2007), is an approach employed in earlier studies investigating the landscape of recombination in non-human primates such as the PanMap (Auton et al. 2012) and Great Ape Recombination Maps (Stevison et al. 2016) projects – which generated fine-scale genetic maps for Western chimpanzees (*Pan troglodytes verus*), Nigerian chimpanzees (*Pan troglodytes ellioti*), bonobos (*Pan paniscus*), and Western gorillas (*Gorilla gorilla gorilla*) – as well as the projects that generated population-scale recombination maps for biomedically-relevant species such as vervet monkeys (*Chlorocebus aethiops sabaeus*; Pfeifer 2020a). The second is the more recently developed software pyrho (Spence and Song 2019) which, unlike LDhat, can explicitly account for the population size change history when performing inference (see “Materials and Methods” section for details).

To assess the performance of these two tools, we simulated a region of 1.6Mb (i.e., the longest accessible intergenic stretch in the aye-aye genome) based on a fixed recombination rate (0.85 cM/Mb; Versoza, Lloret-Villas et al. 2024), mutation rate (0.4 x 10^−8^ and 1.1 x 10^−8^ per base pair per generation; Versoza et al. 2024a), and the recently estimated demographic history for the species consisting of multiple population declines (Terbot et al. 2024), as well as a constant population size for comparison. Our simulations demonstrate that LDhat generally performs well, with estimates falling within the range of the defined recombination rate even under the non-equilibrium demographic model (Figure 2). In contrast, pyrho consistently underestimates recombination rates across all parameter combinations, despite utilizing the defined demographic model during inference. Taken together, these results suggest that LDhat is the superior estimator; additionally, they highlight that the LDhat estimates are themselves relatively robust to the underlying demographic history characterizing aye-ayes.

**Figure 2:**
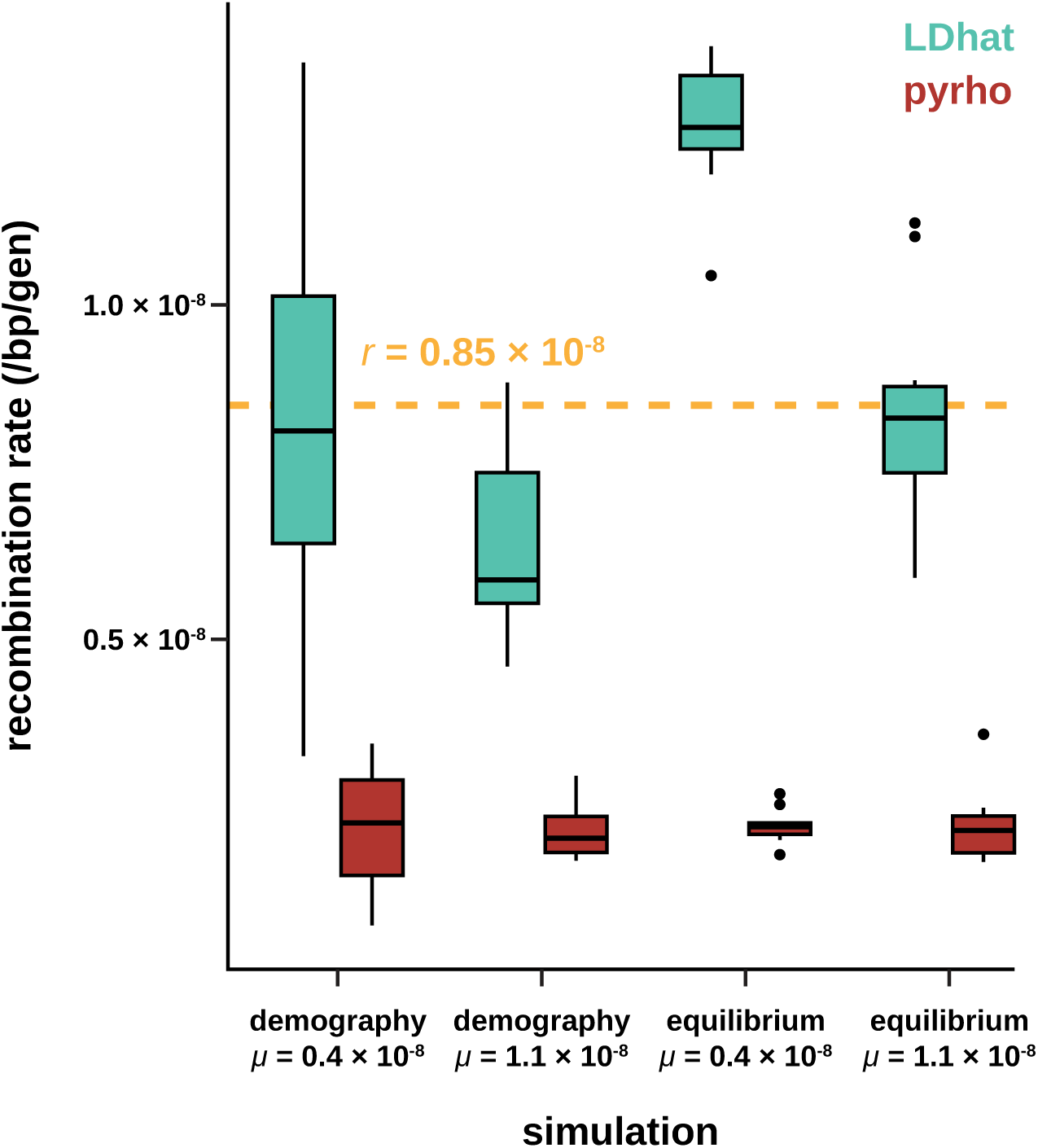
Performance of two common estimators of recombination – the demography-unaware estimator LDhat (shown in teal) and the demography-aware estimator pyrho (shown in red) – across varying mutation rates (*µ* = 0.4 x 10^−8^ and 1.1 x 10^−8^ per base pair per generation [/bp/gen]; Versoza et al. 2024a) and demographic histories, including the demographic history recently estimated by Terbot et al. (2024) for the species consisting of multiple population declines (demography) as well as a constant population size (equilibrium) for comparison. The yellow dashed line depicts the recombination rate that was used in the simulations (i.e., 0.85 cM/Mb; Versoza, Lloret-Villas et al. 2024).

Assuming an ancestral population size of ∼11,750 diploid genomes as recently inferred in the demographic model of Terbot et al. (2024), we thus converted the population-scaled recombination rate estimates inferred using LDhat to per-generation recombination rate estimates, yielding an average genome-wide recombination rate of 1.04 x 10^−9^ per base pair at the 1Mb-scale (Supplementary Figure S15) – about an order of magnitude lower than the average rate reported for anthropoid apes (∼10^−8^ recombination events per base pair per generation, or ∼1 cM/Mb, for humans and ∼1.2 cM/Mb for bonobos, chimpanzees, and gorillas; Kong et al. 2002; Auton et al. 2012; Stevison et al. 2016). This observation of a notable reduction of recombination rates in aye-ayes compared to humans and other haplorrhines is consistent with pedigree-based estimates of sex-specific crossover rates being considerably lower in aye-ayes than in the great apes (Versoza, Lloret-Villas et al. 2024).

However, despite the reduction in overall rate, aye-ayes exhibit a landscape of recombination similar to those of other primates (Auton et al. 2012; Stevison et al. 2016; Pfeifer 2020a; Wall et al. 2022; Versoza, Weiss, et al. 2024); for example, recombination rates are generally elevated towards the telomeric ends and depressed within centromeric and pericentromeric regions of each autosomal scaffold (see Figure 3 for genome-wide recombination rates and Supplementary Figures S16-S29 for the fine-scale variation in recombination rates across each individual autosomal scaffold). Moreover, in aye-ayes, about 80% of recombination occurs in approximately 8% of the genome (Figure 4) – the same fraction than in human individuals of European ancestry (Auton et al. 2012) – potentially hinting at similarities in the concentration of hotspots across the genome.

**Figure 3:**
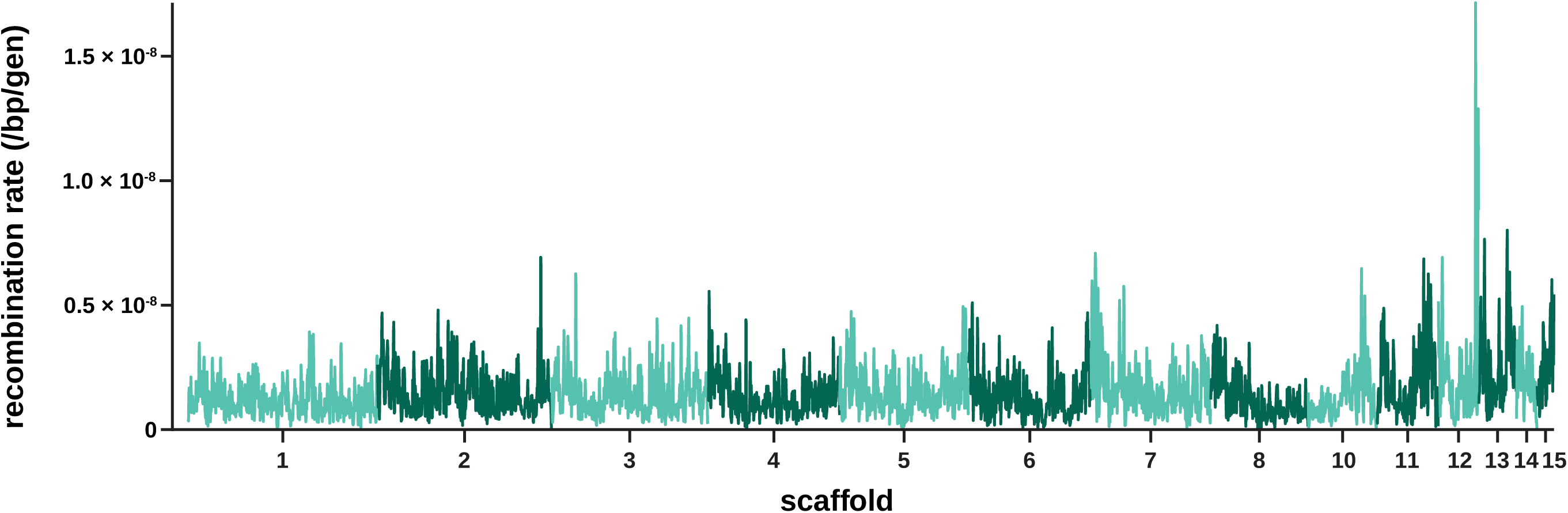
Genome-wide per-base per-generation (/bp/gen) recombination rates for genomic windows of size 1Mb, with a 500kb step size (and see Supplementary Figures S16-S29 for the recombination rate heterogeneity across each individual autosomal scaffold).

**Figure 4:**
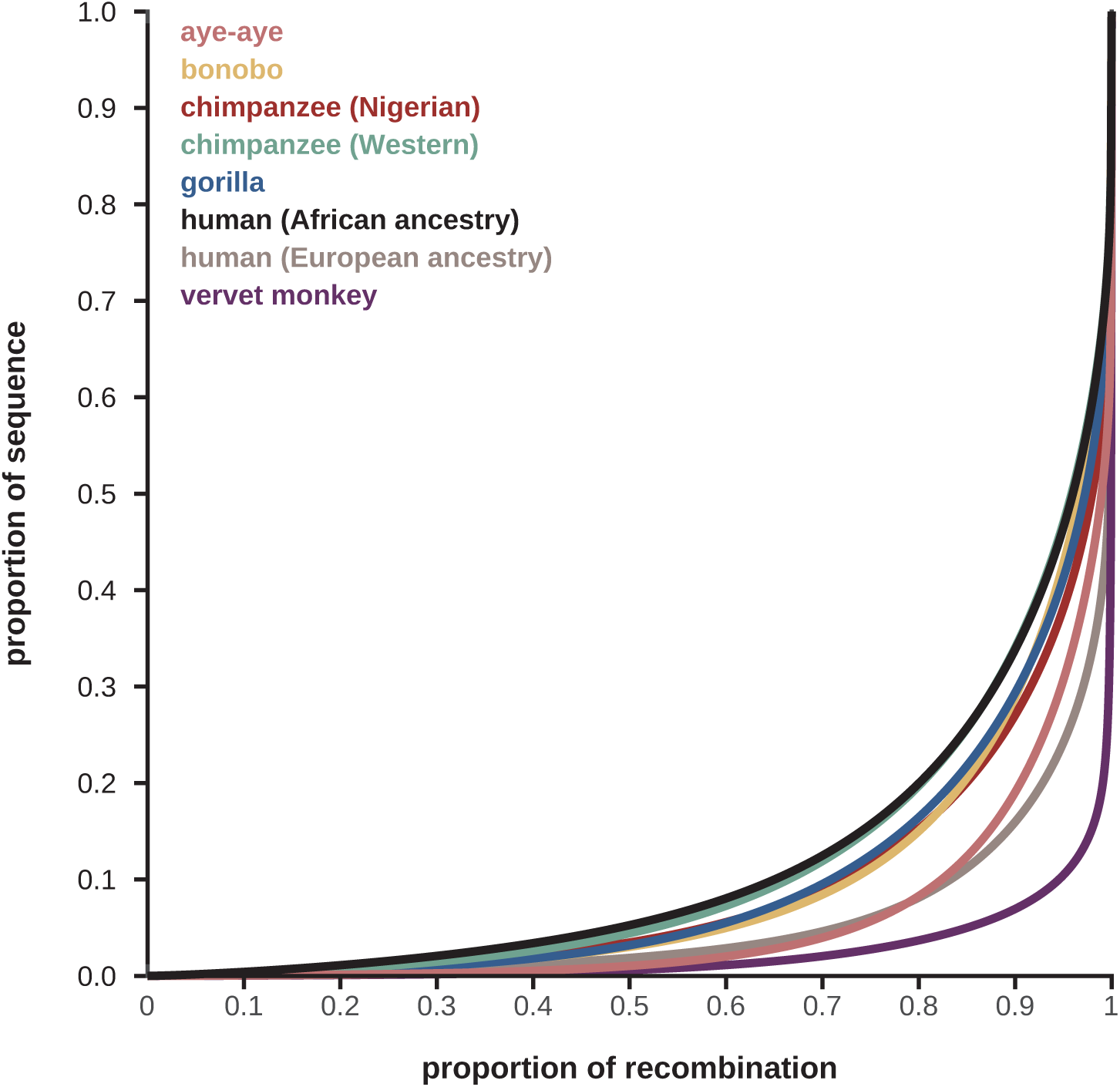
Comparison of the genome-wide distribution of fine-scale recombination rates in aye-ayes (shown in pink) with those of different haplorrhines (with humans of African ancestry shown in black and of European ancestry in beige, chimpanzees originating from Western populations in teal [Auton et al. 2012] and from Nigerian populations in red, bonobos in yellow, gorillas originating from Western populations in blue [Stevison et al. 2016], and vervet monkeys in purple [Pfeifer 2020a]). The figure was adapted from Pfeifer 2020a to include aye-ayes.

### Correlations between fine-scale rates of recombination with genomic features

In order to gain a better understanding of the evolution of the recombination landscape in aye-ayes, we studied the impact of several genomic features on scale-specific recombination rates. To this end, we calculated nucleotide diversity and divergence based on the aye-aye population genomic data and the 447-way mammalian multiple species alignment as noted above, as well as GC-content (as a measure for genome composition) and exon-content (as a proxy for evolutionary constraint) based on the annotated aye-aye assembly in 1kb-regions along the genome. We applied a discrete wavelet transformation in order to obtain information on the heterogeneity in each feature, with detail coefficients providing scale-specific information at a range of (2^n^) scales. After transformation, we performed a linear model analysis of these detail coefficients to study the scale-specific relationships between the heterogeneity in each genomic feature and recombination rate.

Figure 5a provides the detail coefficients for each genomic feature (diagonal plots) as well as their pairwise correlations (off-diagonal plots) at scales ranging from 2^1^ to 2^17^, and Figure 5b the corresponding linear model analysis of the detail coefficients for the longest autosomal scaffold as an example (for all other autosomal scaffolds, see Supplementary Figures S30-S42). Similar to haplorrhines (Spencer et al. 2016; Pfeifer 2020a), aye-ayes exhibit the highest level of heterogeneity in nucleotide diversity and neutral divergence at the finest (2kb) scale. In contrast, the largest heterogeneity in recombination rate occurs over scales of 2-8kb, in the same range previously observed in vervet monkeys (2kb; Pfeifer 2020a) and humans (8kb; Spencer et al. 2006), and similar to the heterogeneity observed in exon-content (4-8kb). Due to the organization of primate genomes into GC-rich and GC-poor isochores (Costantini et al. 2009), base composition displays a concave distribution, with the highest heterogeneities observed at both the fine (2-8kb) and broad (>1Mb) scales. Focusing on the pairwise correlations between the detail coefficients at the fine (2-8kb) scale, nucleotide diversity is significantly positively correlated with both neutral divergence and GC-content, as expected given that the rate of mutation, which jointly impacts diversity and divergence, varies depending on the local base composition in the genome (Figure 1c, and see review of Hodgkinson and Eyre-Walker 2011). The rates of divergence are also significantly negatively correlated with exon-content at the fine to intermediate scales, as anticipated from evolutionary constraint to maintain proper gene function, thereby subjecting these regions to purifying selection (see reviews of Charlesworth and Jensen 2021, 2022). In addition to mutation, and similar to other primates (Spencer et al. 2006; Auton et al. 2012; Pfeifer and Jensen 2016; Stevison et al. 2016), GC-rich genomic regions are also associated with higher rates of recombination in aye-ayes. Contributing to this positive correlation at the fine-scale is GC-biased gene conversion, an evolutionary process associated with meiotic recombination that elevates the GC-content of a region through the preferential transmission of GC over AT alleles (Duret and Galtier 2009), thus leading to a higher GC-content in regions of frequent recombination (i.e., recombination hotspots). Additionally, in regions of high recombination, the effects of selection at linked sites (e.g., background selection and selective sweeps) will be reduced, allowing more genetic diversity to persist in close proximity (Maynard Smith and Haigh 1974; Begun and Aquadro 1992; Charlesworth et al. 1993). However, recombination hotspots are highly localized (within 1-2kb regions; Baudat et al. 2010; Myers et al. 2010; Parvanov et al. 2010) and often flanked by regions of low recombination which, in turn, extend genetic hitchhiking effects, thus reducing nucleotide diversity at intermediate (10s to 100s of kb) scales (Maynard Smith and Haigh 1974; Begun and Aquadro 1992; Charlesworth et al. 1993).

**Figure 5:**
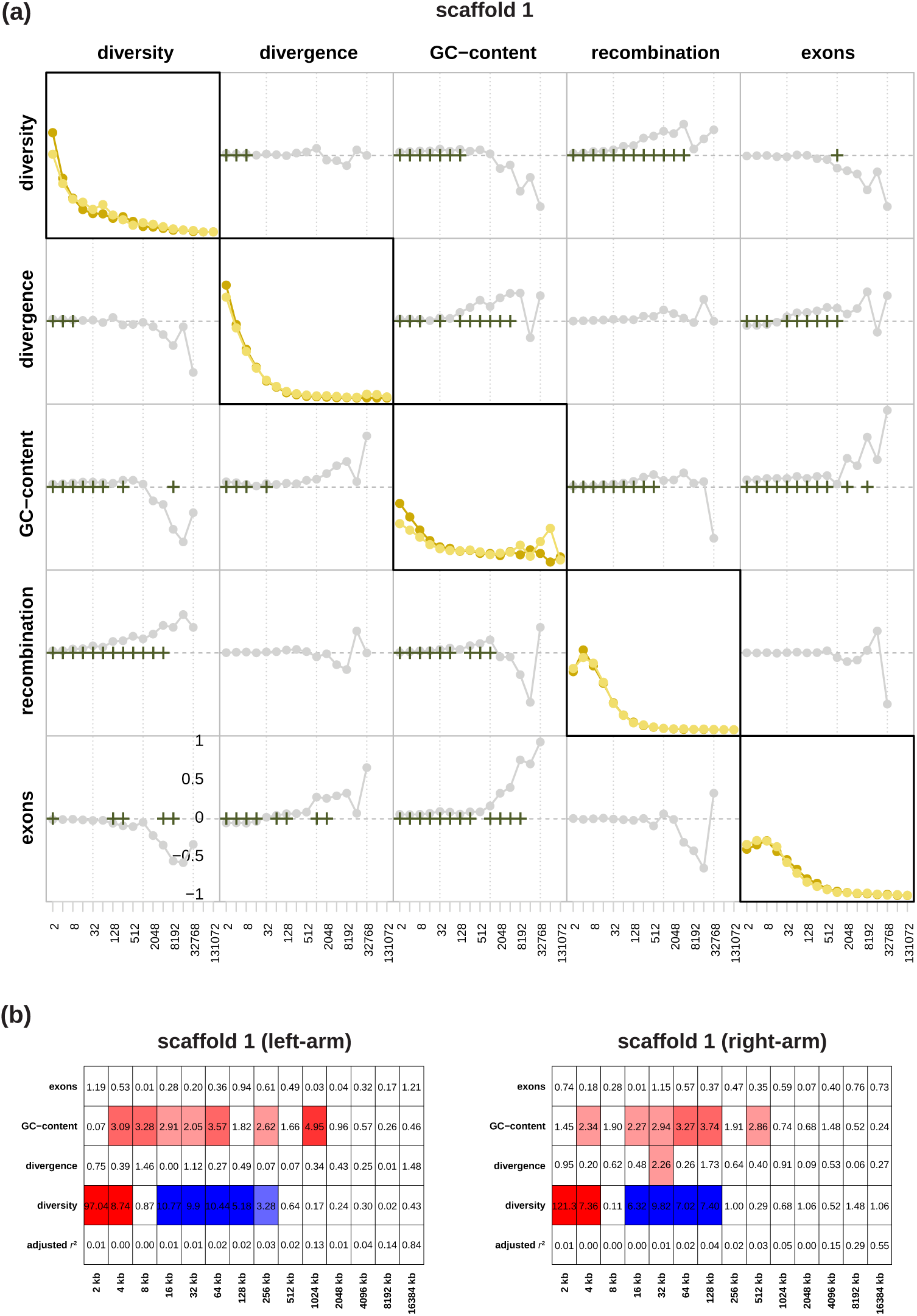
(a) The detail coefficients of each genomic feature (diagonal plots) on the left and right arms of scaffold 1 (shown in dark and light yellow, respectively) as well as their pairwise correlations based on Kendall’s rank correlation (off-diagonal plots with the bottom left showing the left-arm and the top right showing the right-arm) at a range of (2^n^) scales. Correlations significant at the 1%-level under a two-tailed test are highlighted by crosses. (b) Linear model analysis of the detail coefficients. Red and blue coloring indicate significant positive and negative relationships under a two-sided *t*-test, with the color intensity being proportional to the significance level. Adjusted *r*^2^ specifies the proportion of heterogeneity that can be explained by the linear model.

## Concluding thoughts

In this study, we have characterized the underlying heterogeneity in mutation and recombination rates across the genome of aye-ayes. We found that mutation rates in this species are lower than in other primates, which is in agreement with previous studies showing lower mutation rates in prosimians (Tran and Pfeifer 2018; Chintalapati and Moorjani 2020). Notably, this indirect divergence-based estimate supports the recent pedigree-based estimate of 0.4 x 10^−8^ per base pair per generation characteristic of younger parents (Versoza et al. 2024a), suggesting a relatively young long-term reproductive age in the wild, as might be expected from previous studies of the life history and socioecology of the species (Ross 2003). This rate also implies a split time of ∼54 mya, consistent with the earliest primates in the fossil record, as opposed to the much older and difficult to reconcile split times previously proposed. We similarly found a notable reduction of recombination rate in aye-ayes compared to the great apes (Auton et al. 2012; Stevison et al. 2016), despite overall similarities in the recombination landscape, including the concentration of hotspots across the genome. Given the recently reported enrichment of crossover events in regions harboring predicted great ape PRDM9 binding motifs – a zinc-finger protein controlling the activation of hotspots in primates – in pedigreed aye-aye individuals (Versoza, Lloret-Villas et al. 2024), the future characterization of hotspots in the species should thus be of great interest to the comparative primate genomics community.

With rate maps available in only a limited number of species, it is common practice to use a single, species-averaged rate for both mutation and recombination when modelling population genetic processes. However, failing to account for the underlying heterogeneity in mutation and recombination rates has been shown to potentially bias the inference of both population history as well as the distribution of fitness effects (e.g., Soni et al. 2023, 2024a). Thus, the rate maps provided here will facilitate more robust inference of population genetic processes in the highly endangered aye-aye specifically, as well as in evolutionary models of primate evolution more broadly.

## Materials and Methods

### Updating the aye-aye genome in the 447-way mammalian multiple species alignment

We obtained the 447-way multiple species alignment, consisting of the combined mammalian multiple species alignment of the Zoonomia Consortium (2020) and the primate multiple species alignment of Kuderna et al. (2024), from https://cglgenomics.ucsc.edu/november-2023-nature-zoonomia-with-expanded-primates-alignment/ and removed the outdated aye-aye genome assembly using the *halRemoveGenome* function implemented in HAL v.2.2 (Hickey et al. 2013). Next, we added the current NCBI reference genome for the species – that is, the high-quality, fully annotated aye-aye assembly of Versoza and Pfeifer (2024) (DMad_hybrid; GenBank accession number: JBFSEQ000000000) – to the alignment, by first extracting the ancestral genomes PrimatesAnc005 and PrimatesAnc011 from the 447-way alignment using HAL’s *hal2fasta* function, and then aligning these ancestral genomes with the new aye-aye genome in Cactus v.2.9.2 (Armstrong et al. 2020) using the branch lengths previously inferred in the 447-way alignment. Finally, we attached this alignment back into the 447-way alignment using HAL’s *halReplaceGenome* function.

### Inferring fine-scale rates of neutral divergence and mutation

To infer fine-scale rates of neutral divergence and mutation, we first used the *halSummarizeMutations* function implemented in HAL v.2.2 (Hickey et al. 2013) to retrieve ‘point mutations’ along the aye-aye branch (i.e., substitutions between the aye-aye and PrimateAnc005), thereby masking any sites within 10kb of functional regions to avoid the potentially confounding effects of selection. From these point mutations, we then removed all sites associated with segregating polymorphisms in the species, resulting in a final dataset from which we calculated neutral divergence by dividing the number of divergent sites by the number of accessible sites in any given genomic window (Soni et al. 2024b). Specifically, divergence was estimated genome-wide, as well as in windows of size 1kb, 10kb, 100kb and 1Mb using a sliding window approach with a step size of 1kb, 5kb, 50kb, and 500kb, respectively. To obtain mutation rates for each genomic window, we divided by the divergence time in generations, using divergence times of 54.9 mya and 74.7 mya (Horvarth et al. 2008) and generation times of 3 years and 5 years (Ross 2003; Louis et al. 2020) for comparison.

### Inferring fine-scale rates of recombination

We utilized two different approaches to infer fine-scale rates of recombination – the demography-unaware estimator LDhat (McVean et al. 2002, 2004; Auton and McVean 2007) and the demography-aware estimator pyrho (Spence and Song 2019) – both of which rely on patterns of LD observed in sequencing data to estimate recombination rates. To this end, we took advantage of a recently generated population genomic dataset of unrelated individuals (Soni et al. 2024b) for which we implemented a set of stringent filter criteria (supplementing the standard quality control practices applied in the previous study as described in Pfeifer 2017) to eliminate spurious single nucleotide polymorphisms (SNPs) that may lead to artefactual breaks in patterns of LD. Specifically, following the guidelines described in earlier studies investigating the landscape of recombination in non-human primates (Auton et al. 2012; Stevison et al. 2016; Pfeifer 2020a), we removed both SNP clusters – defined here as three or more SNPs within a 10bp window (calculated using the Genome Analysis Toolkit [GATK] v.4.2.6.1 *VariantFiltration* function together with the parameters ‘ ––cluster-size 3 ‘ and ‘ ––cluster-window-size 10 ‘; van der Auwera and O’Connor 2020) – as well as SNPs exhibiting an excess of heterozygosity – defined here as sites with a Hardy-Weinberg equilibrium *p*-value of < 0.01 (calculated using the ‘ ––hardy ‘ option in VCFtools v.0.1.14; Danecek et al. 2011) – from the published dataset. Additionally, we excluded all SNPs located within regions blacklisted by the ENCODE Project Consortium (2012) (i.e., within regions prone to artifacts in high-throughput sequencing experiments) by lifting the data between the aye-aye (DMad_hybrid) genome assembly and the human (hg38) genome assembly using the UCSC liftOver tool (Raney et al. 2024). The resulting high-quality, population-level dataset, consisting of 3,454,304 biallelic autosomal SNPs (transition-transversion ratio: 2.53), was then used as input for the recombination rate estimators LDhat (McVean et al. 2002, 2004; Auton and McVean 2007) and pyrho (Spence and Song 2019).

> *<u>LDhat</u>*: Following previous work in catarrhines (Auton et al. 2012; Stevison et al. 2016; Pfeifer 2020a), we estimated the population recombination rate using LDhat v.2.2 (McVean et al. 2002, 2004; Auton and McVean 2007). In brief, we first divided the high-quality population-level dataset into 4,000-SNP regions with a 200-SNP overlap between adjacent regions, and then ran the *interval* function of LDhat with a block penalty of 5 (’ –bpen 5 ‘) for 60 million iterations (’ –its 60000000 ‘) using a sampling scheme of 40,000 iterations (’ –samp 40000 ‘). Afterward, we used LDhat’s *stat* function to discard the burn-in – defined here as the first 20 million iterations (’ –burn 500 ‘) of the Monte Carlo Markov Chain – and combined the region-based recombination rate estimates at the midpoint of the overlap. In keeping with previous best practices, we checked for regions with recombination rate estimates of > 100 between adjacent SNPs as well as gaps > 50 kb in the genome assembly that might spuriously interrupt patterns of LD, but no such regions were identified. Lastly, as LDhat estimates the population recombination rate ρ = 4 *N_e_ r*, where *N_e_* is the effective population size and *r* is the per-generation recombination rate, we used the ancestral population size inferred in the demographic model of Terbot et al. (2024) (i.e., ∼11,750 diploid genomes) to convert ρ to *r*.

> *<u>pyrho</u>*: Following the recommendations of the developers (Spence and Song 2019), we estimated the per-generation recombination rate *r* using pyrho v.0.1.7. In brief, we first generated a likelihood lookup table using pyrho’s *make_table* function, taking into account the population size change history previously inferred by Terbot et al. (2024) (’ ––popsizes 2570,2944.784,3374.224,3866.288,4430.111,5076.157,5816.415,6585,23389 ––epochtimes 1,2,3,4,5,6,7,1133 ‘), and then ran the *hyperparam* function with the species-specific mutation rate estimated by Versoza et al. (2024a) for individuals reproducing at a young age (’ ––mu 0.4e-8 ‘), as likely the case in the wild (Ross 2003), to determine the optimal parameter settings for window size and block penalty. We then used pyrho’s *optimize* function with the recommended window size of 30 (’ ––windowsize 30 ‘) and block penalty of 45 (’ ––blockpenalty 45 ‘) to estimate per-generation recombination rates across the genome.

### Assessing the performance of recombination rate estimators using simulations

To compare the performance of the demography-unaware recombination rate estimator LDhat with the demography-aware estimator pyrho, we used msprime v.1.3.2 (Baumdicker et al. 2022) to simulate 10 replicates of a 1.6Mb region (i.e., the longest uninterrupted accessible intergenic region in the aye-aye genome) with multiple parameter combinations. Specifically, to test the robustness of both tools with regards to the underlying demographic history, we implemented two models in our simulations: (1) the bottleneck-decline model from Terbot et al. (2024) and (2) a constant equilibrium model. Moreover, in addition to the species-specific average mutation rate recently estimated from a 14-individual three-generation pedigree in Versoza et al. (2024a) (1.1 x 10^−8^ per base pair per generation), we also considered the lowest reported pedigree estimate (0.4 x 10^−8^ per base pair per generation) in our models to account for individuals potentially reproducing at a young age in the wild. Finally, we used the coarse-scale recombination rate estimate from pedigreed individuals (0.85 cM/Mb) reported in Versoza, Lloret-Villas et al. (2024) in all models.

### Assessing the correlation of fine-scale rates of recombination with genomic features

Following previous work in humans (Spencer et al. 2006), we first calculated nucleotide diversity and divergence based on the aye-aye population genomic data and the 447-way mammalian multiple species alignment as noted above, as well as GC-content (as a measure of base composition) and exon-content (as a proxy for evolutionary constraint) based on the annotated aye-aye (DMad_hybrid) genome assembly (GenBank accession number: JBFSEQ000000000; Versoza and Pfeifer 2024) in 1kb windows along the 14 autosomal scaffolds (i.e., scaffolds 1-8 and 10-15), and then applied a discrete wavelet transformation using the *Rwave* and *wavethresh* packages implemented in R v.4.2.2 to obtain information on the heterogeneity in each genomic feature at varying scales. To study scale-specific correlations, we additionally performed a linear model analysis on the log-transformed recombination, nucleotide diversity, and divergence rates.

## Acknowledgements

We would like to thank the Duke Lemur Center for providing the aye-aye samples used in this study, and members of the Jensen Lab and Pfeifer Lab for helpful discussion. Computations were performed on the Sol supercomputer at Arizona State University (Jennewein et al. 2023) and on the Open Science Grid, which is supported by the National Science Foundation and the U.S. Department of Energy’s Office of Science. This is Duke Lemur Center publication # XXXX.

## Funding

This work was supported by the National Institute of General Medical Sciences of the National Institutes of Health under Award Number R35GM151008 to SPP and the National Science Foundation under Award Number DBI-2012668 to the Duke Lemur Center. VS, JT, and JDJ were supported by National Institutes of Health Award Number R35GM139383 to JDJ. CJV was supported by the National Science Foundation CAREER Award DEB-2045343 to SPP. The content is solely the responsibility of the authors and does not necessarily represent the official views of the National Institutes of Health or the National Science Foundation.

## Notes

### Competing Interest Statement

The authors have declared no competing interest.

