## Supplementary Materials for "Inferring fine-scale mutation and recombination rate maps in aye-ayes (*Daubentonia madagascariensis*)"

**S1**

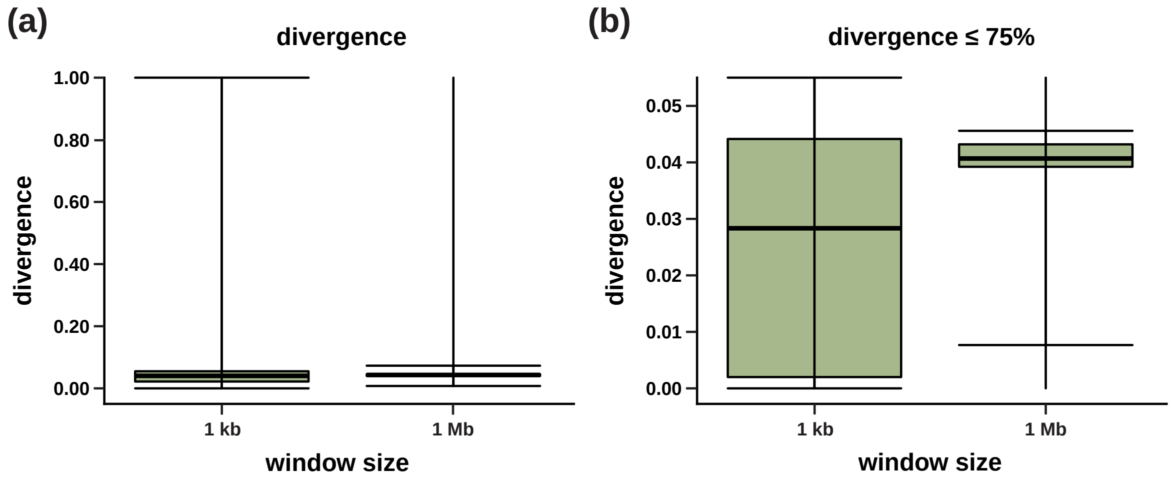


**Supplementary Figure S1:** Distribution of divergence for (a) all 1kb and 1Mb genomic windows and (b) for windows in the lower three quartiles. Bold bars represent mean divergence, with boxes representing the 25% and 75% quartiles. Whiskers represent minimum and maximum values.

**S2**

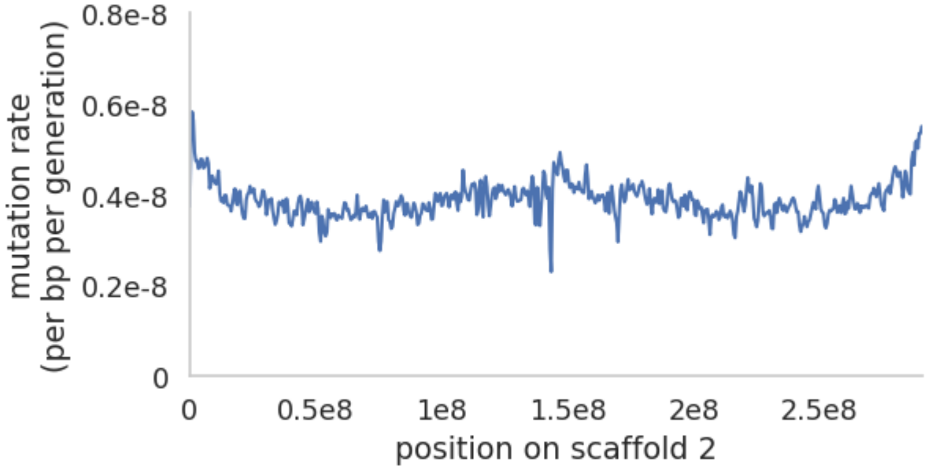


**Supplementary Figure S2:** Fine-scale mutation rates along scaffold 2 for genomic windows of size 1Mb, with a 500kb step size.

**S3**

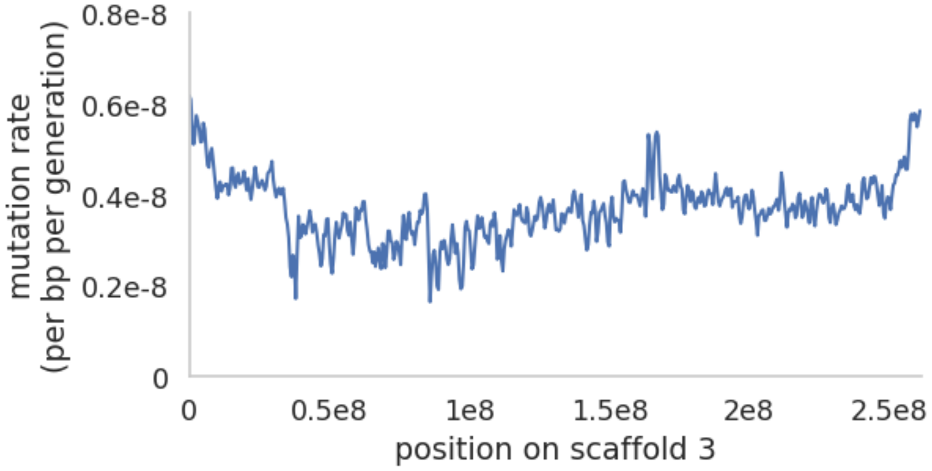


**Supplementary Figure S3:** Fine-scale mutation rates along scaffold 3 for genomic windows of size 1Mb, with a 500kb step size.

**S4**

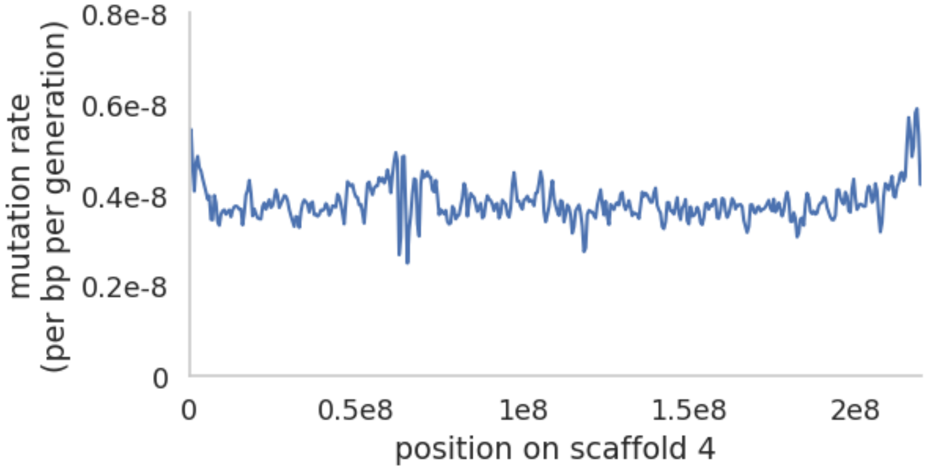


**Supplementary Figure S4:** Fine-scale mutation rates along scaffold 4 for genomic windows of size 1Mb, with a 500kb step size.

**S5**

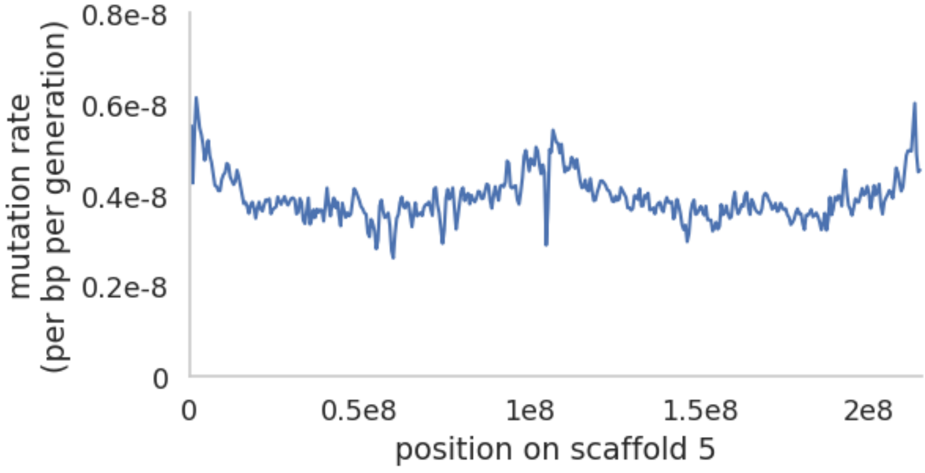


**Supplementary Figure S5:** Fine-scale mutation rates along scaffold 5 for genomic windows of size 1Mb, with a 500kb step size.

**S6**

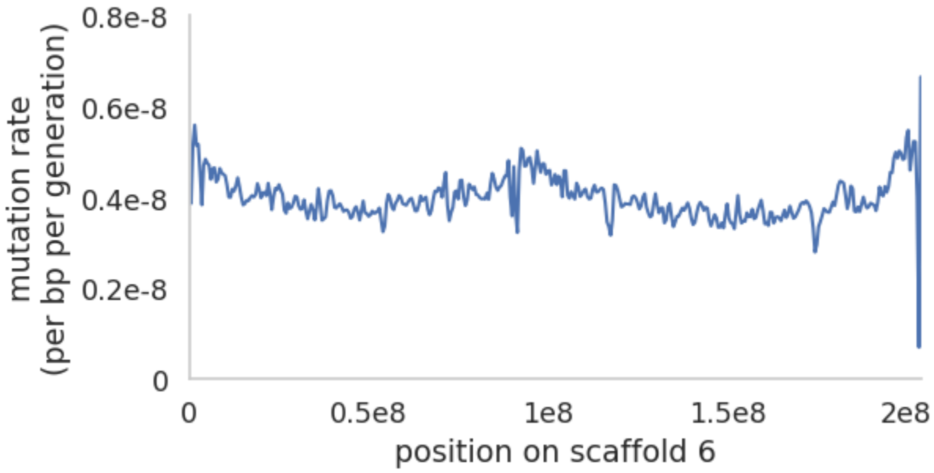


**Supplementary Figure S6:** Fine-scale mutation rates along scaffold 6 for genomic windows of size 1Mb, with a 500kb step size.

**S7**

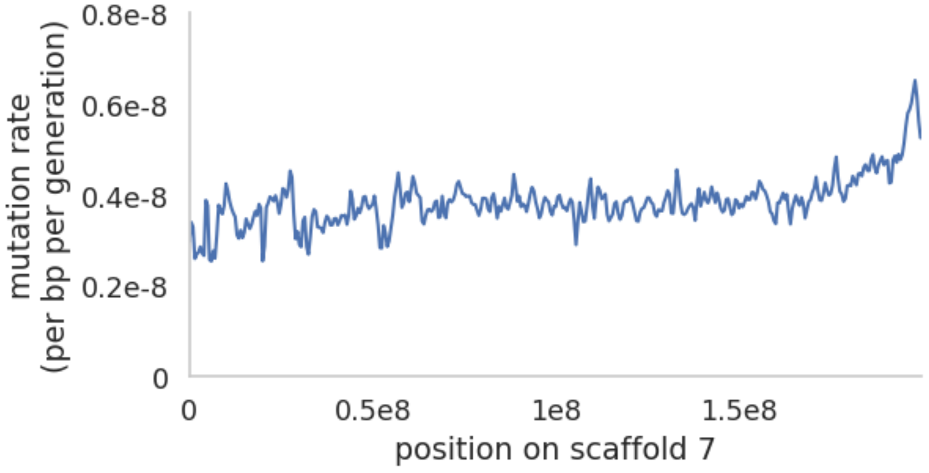


**Supplementary Figure S7:** Fine-scale mutation rates along scaffold 7 for genomic windows of size 1Mb, with a 500kb step size.

**S8**

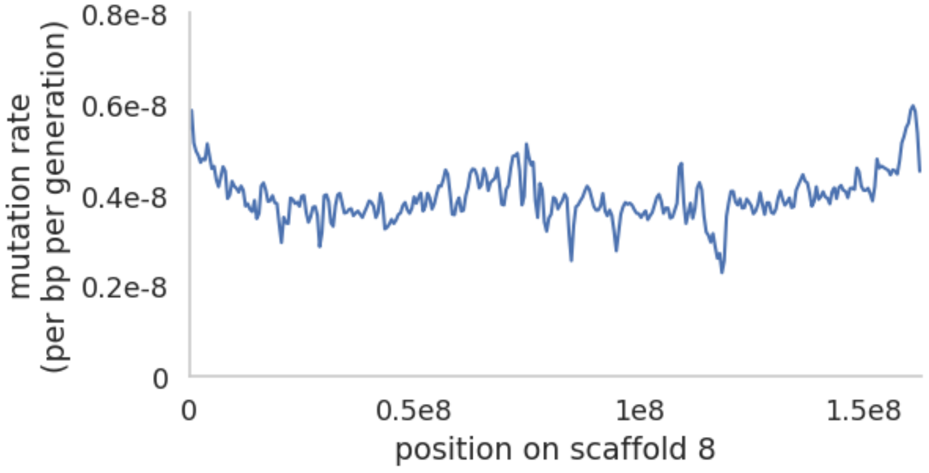


**Supplementary Figure S8:** Fine-scale mutation rates along scaffold 8 for genomic windows of size 1Mb, with a 500kb step size.

**S9**

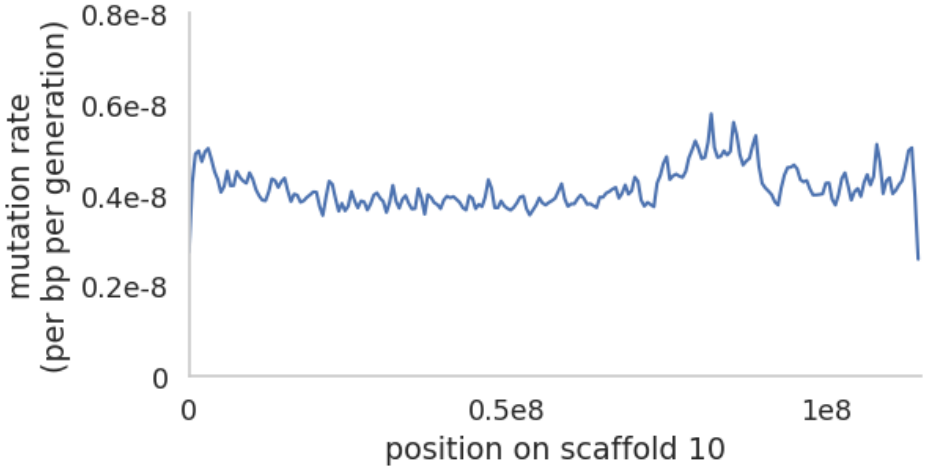


**Supplementary Figure S9:** Fine-scale mutation rates along scaffold 10 for genomic windows of size 1Mb, with a 500kb step size.

**S10**

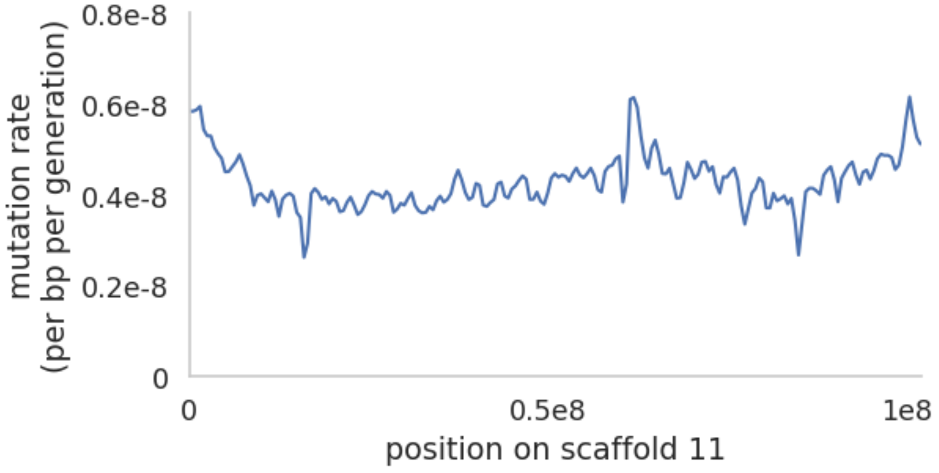


**Supplementary Figure S10:** Fine-scale mutation rates along scaffold 11 for genomic windows of size 1Mb, with a 500kb step size.

**S11**

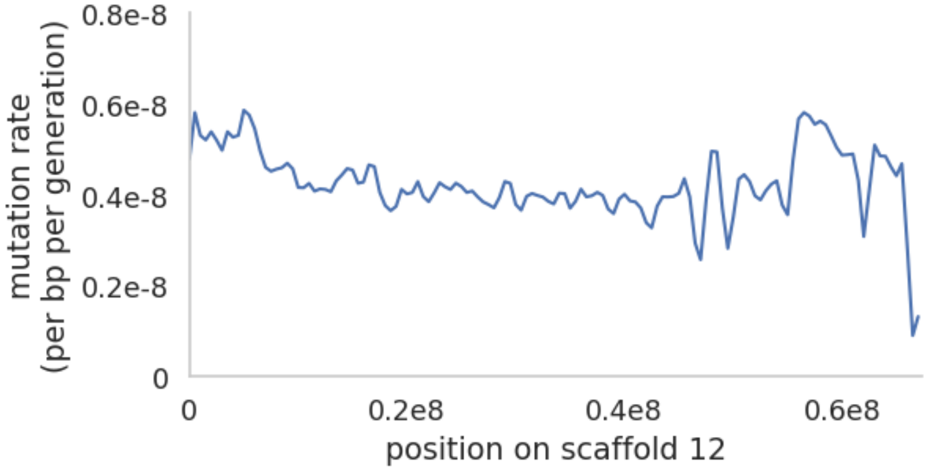


**Supplementary Figure S11:** Fine-scale mutation rates along scaffold 12 for genomic windows of size 1Mb, with a 500kb step size.

**S12**

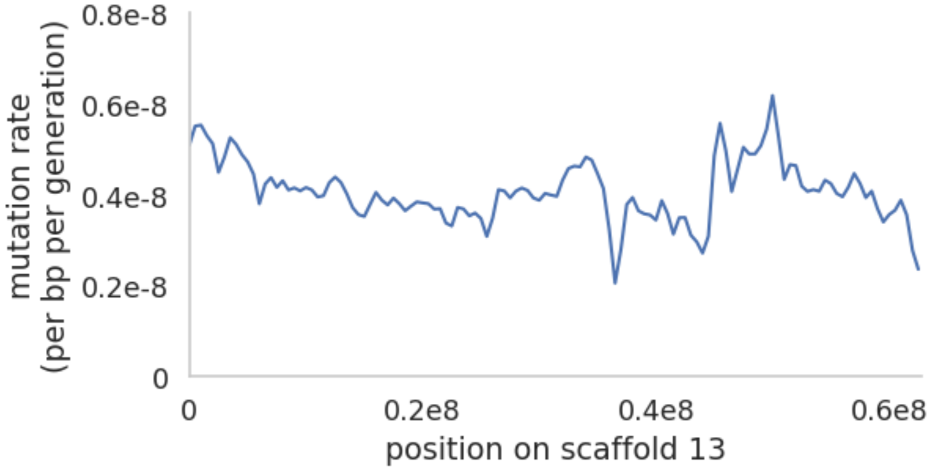


**Supplementary Figure S12:** Fine-scale mutation rates along scaffold 13 for genomic windows of size 1Mb, with a 500kb step size.

**S13**

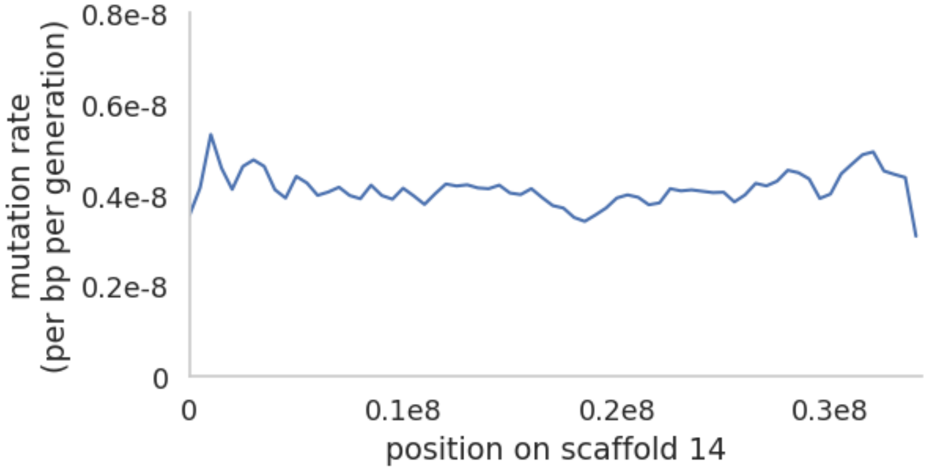


**Supplementary Figure S13:** Fine-scale mutation rates along scaffold 14 for genomic windows of size 1Mb, with a 500kb step size.

**S14**

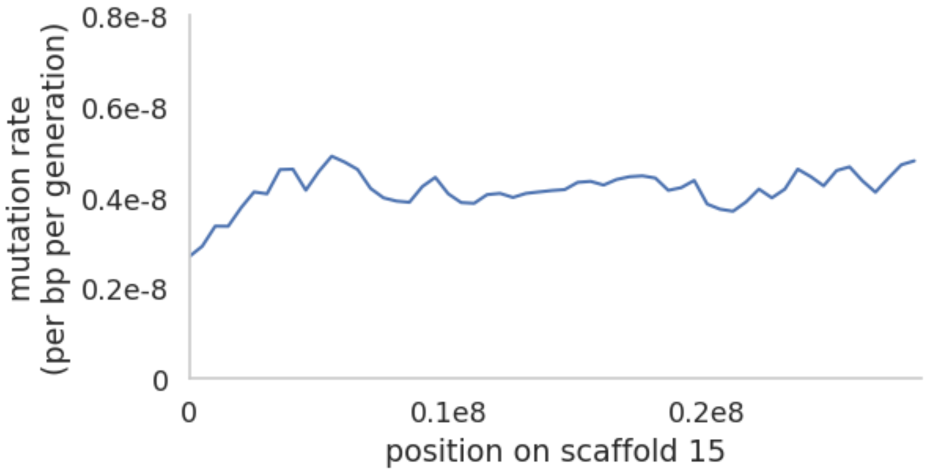


**Supplementary Figure S14:** Fine-scale mutation rates along scaffold 15 for genomic windows of size 1Mb, with a 500kb step size.

**S15**

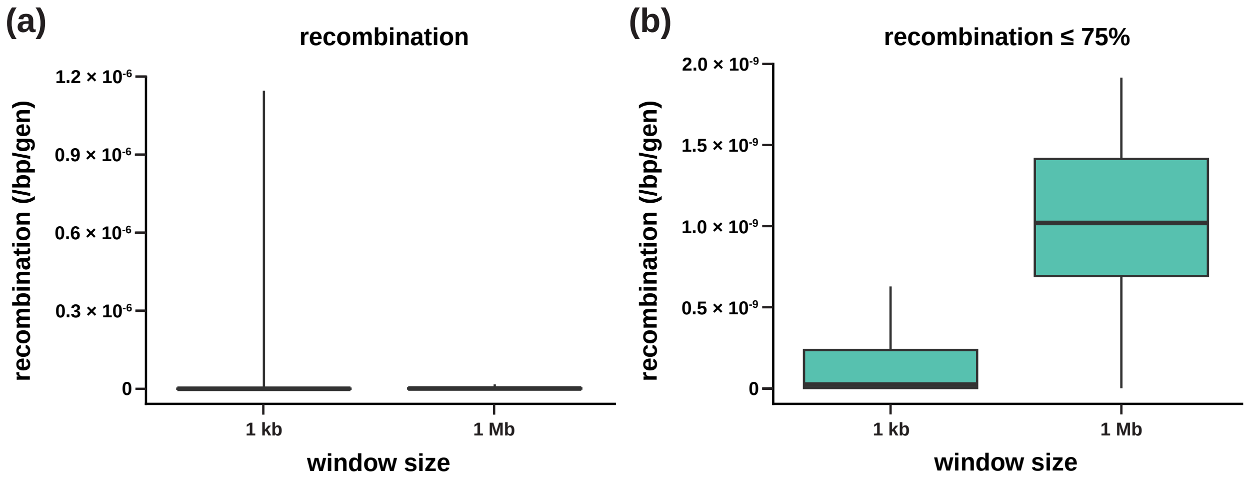


**Supplementary Figure S15**: Distribution of per-base per-generation recombination rates for (a) all 1kb and 1Mb genomic windows and (b) for windows in the lower three quartiles. Bold bars represent mean recombination rates, with boxes representing the 25% and 75% quartiles.

## S16

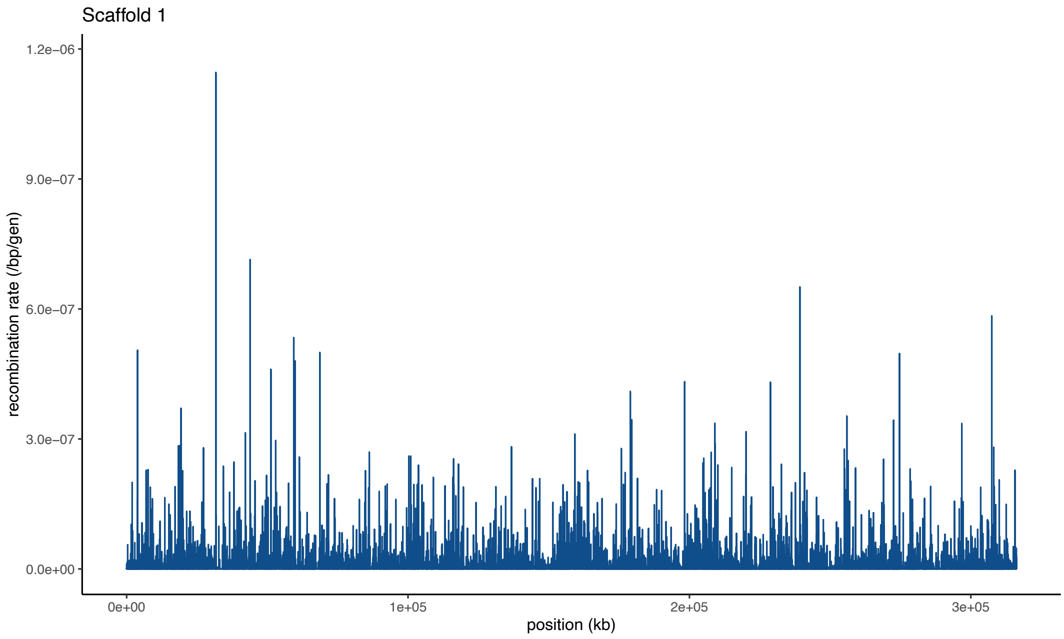


**Supplementary Figure S16:** Fine-scale per-base per-generation (/bp/gen) recombination rates along scaffold 1 for genomic windows of size 1Mb, with a 500kb step size.

**S17**

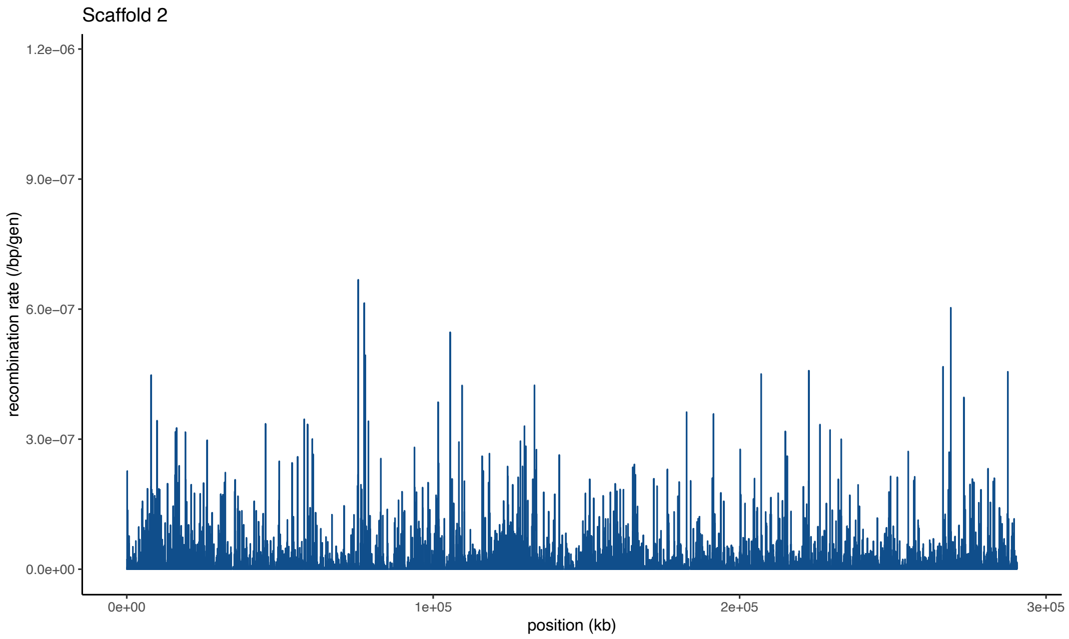


**Supplementary Figure S17:** Fine-scale per-base per-generation (/bp/gen) recombination rates along scaffold 2 for genomic windows of size 1Mb, with a 500kb step size.

## S18

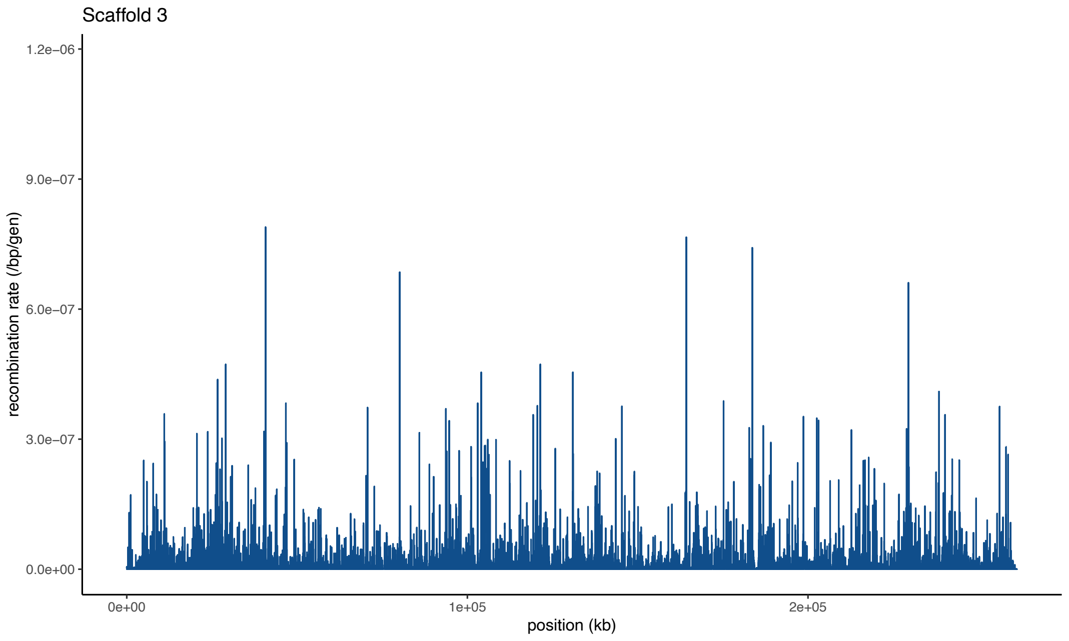


**Supplementary Figure S18:** Fine-scale per-base per-generation (/bp/gen) recombination rates along scaffold 3 for genomic windows of size 1Mb, with a 500kb step size.

**S19**

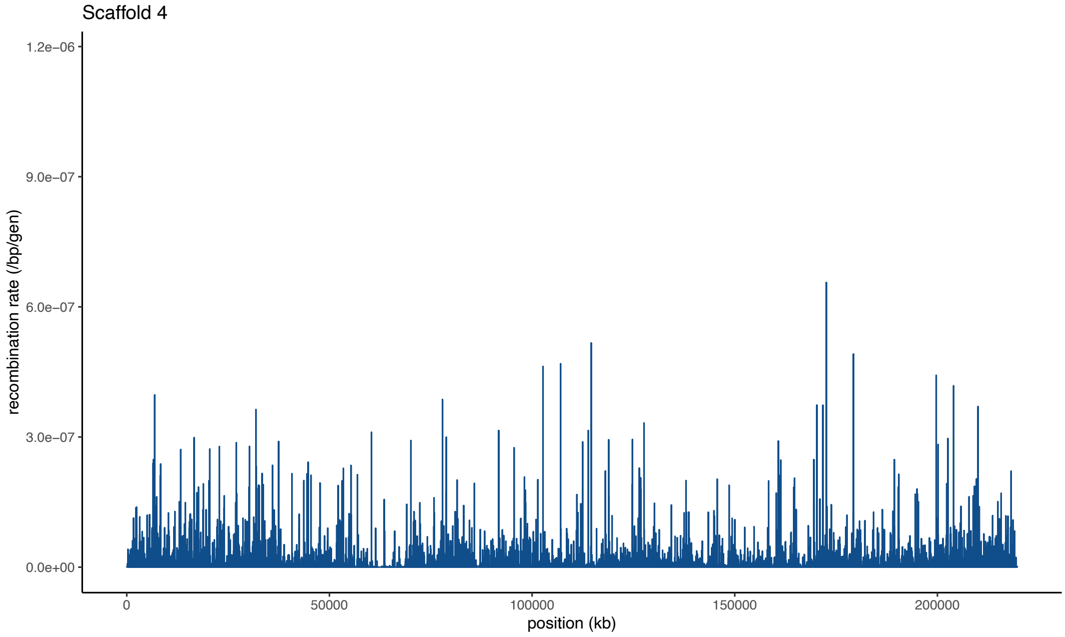


**Supplementary Figure S19:** Fine-scale per-base per-generation (/bp/gen) recombination rates along scaffold 4 for genomic windows of size 1Mb, with a 500kb step size.

**S20**

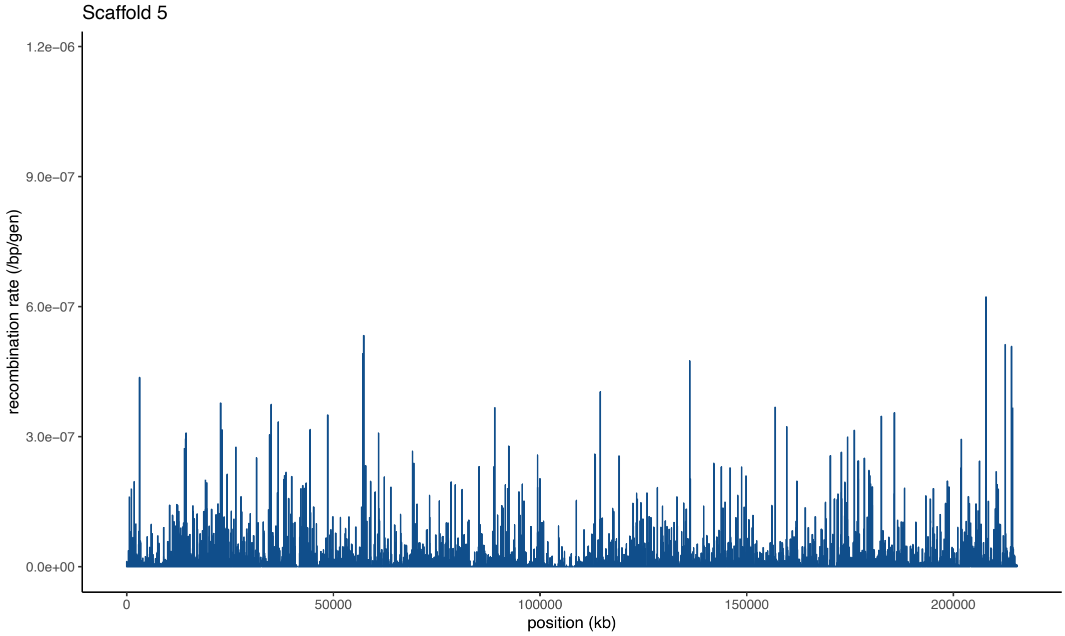


**Supplementary Figure S20:** Fine-scale per-base per-generation (/bp/gen) recombination rates along scaffold 5 for genomic windows of size 1Mb, with a 500kb step size.

**S21**

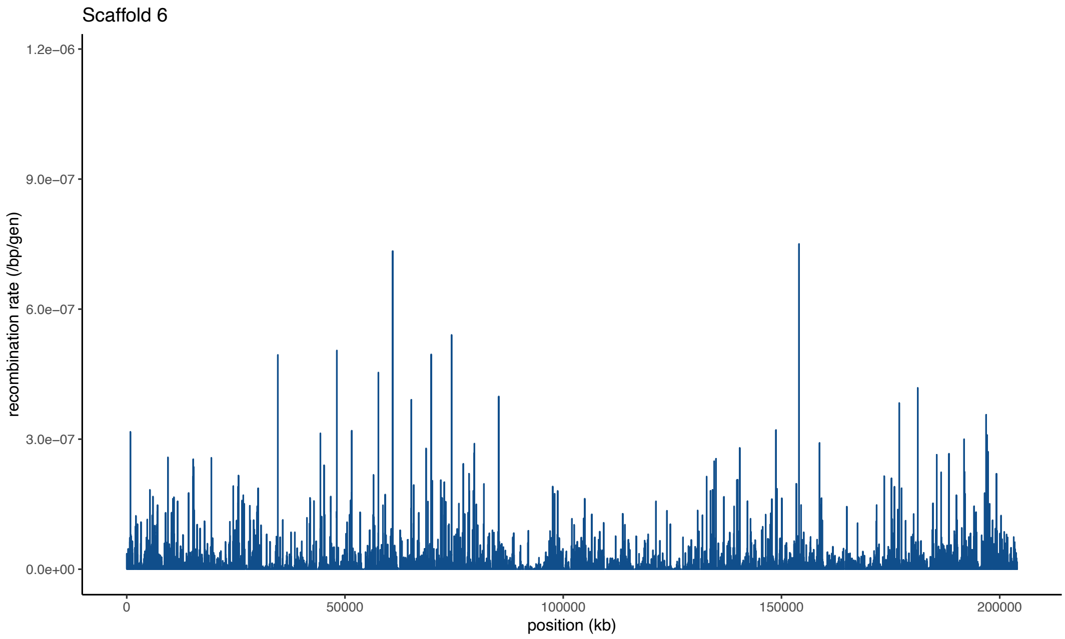


**Supplementary Figure S21:** Fine-scale per-base per-generation (/bp/gen) recombination rates along scaffold 6 for genomic windows of size 1Mb, with a 500kb step size.

**S22**

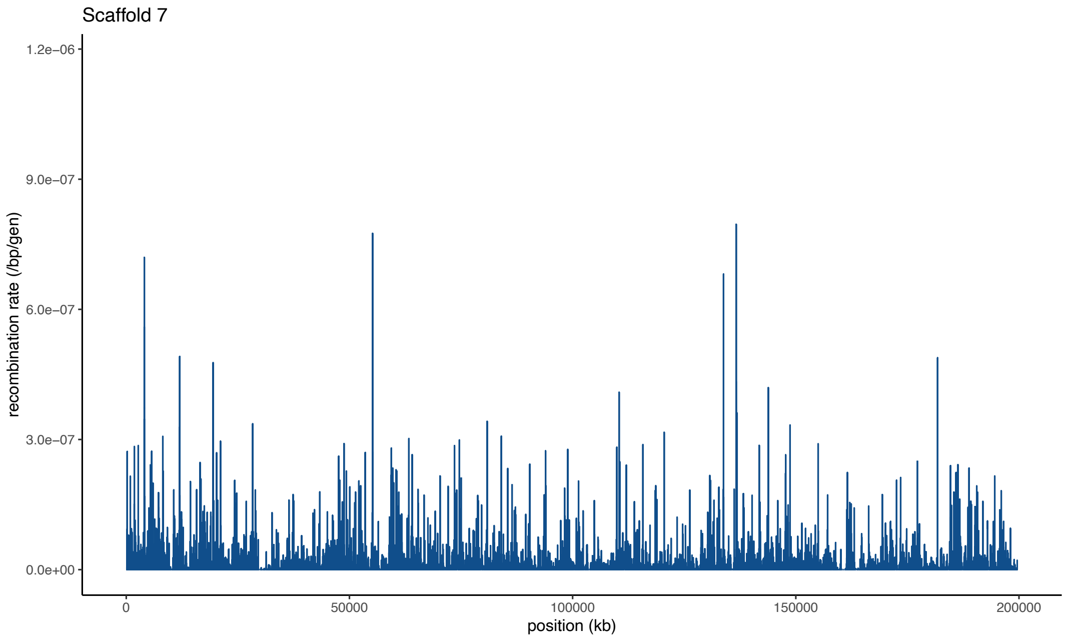


**Supplementary Figure S22:** Fine-scale per-base per-generation (/bp/gen) recombination rates along scaffold 7 for genomic windows of size 1Mb, with a 500kb step size.

**S23**

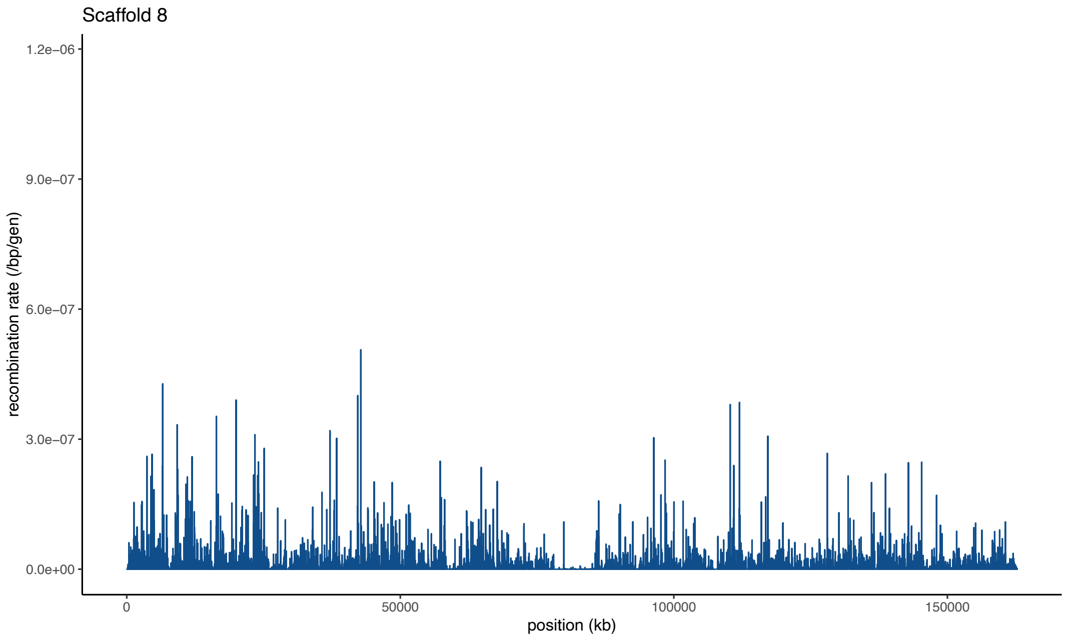


**Supplementary Figure S23:** Fine-scale per-base per-generation (/bp/gen) recombination rates along scaffold 8 for genomic windows of size 1Mb, with a 500kb step size.

**S24**

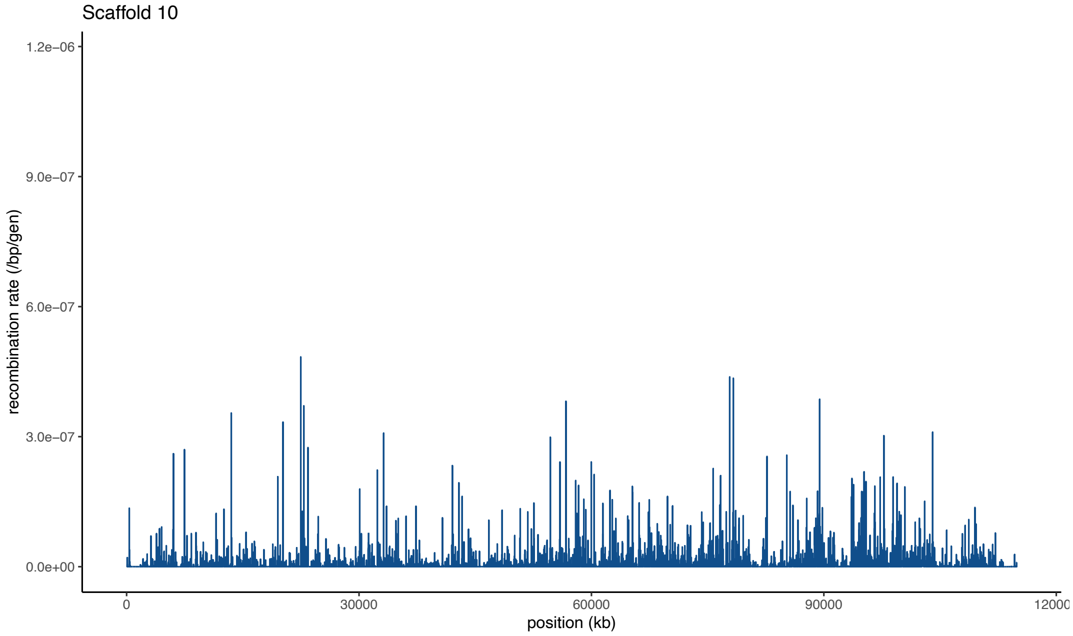


**Supplementary Figure S24:** Fine-scale per-base per-generation (/bp/gen) recombination rates along scaffold 10 for genomic windows of size 1Mb, with a 500kb step size.

**S25**

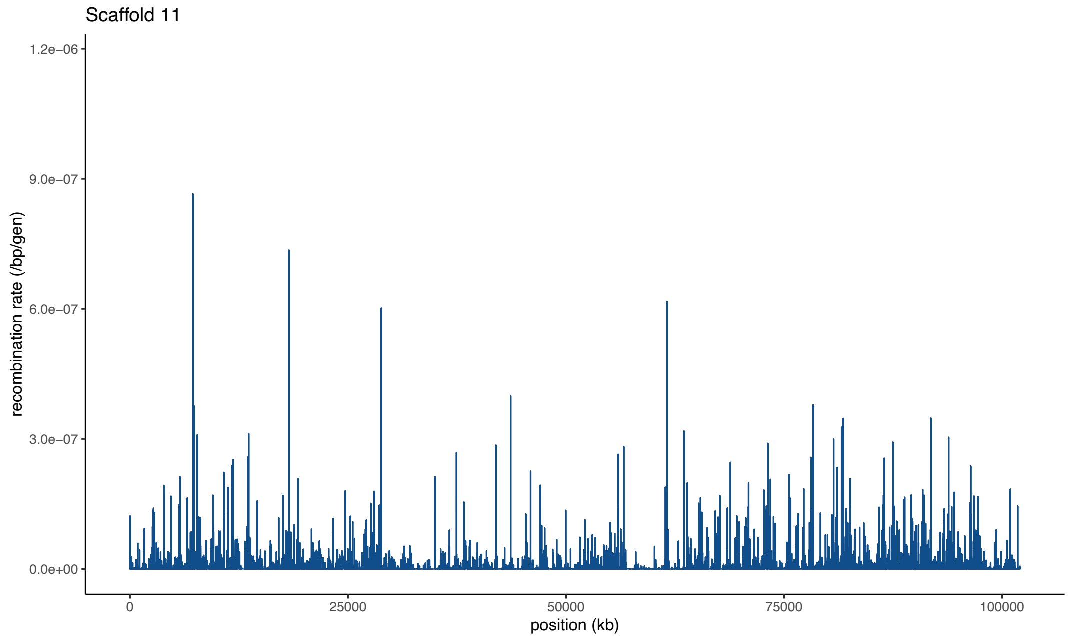


**Supplementary Figure S25:** Fine-scale per-base per-generation (/bp/gen) recombination rates along scaffold 11 for genomic windows of size 1Mb, with a 500kb step size.

**S26**

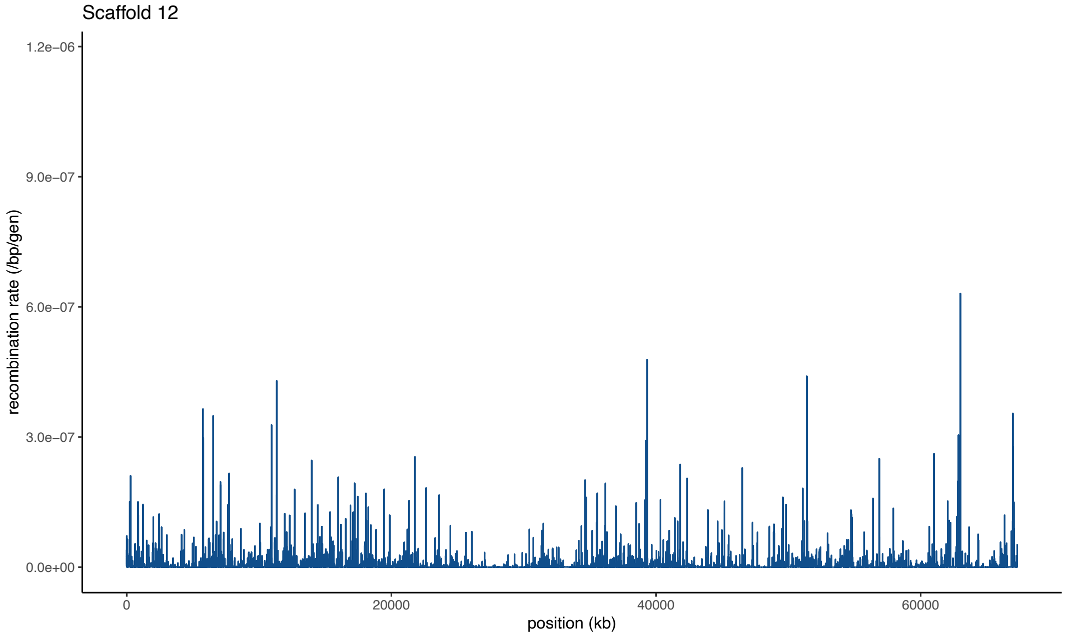


**Supplementary Figure S26:** Fine-scale per-base per-generation (/bp/gen) recombination rates along scaffold 12 for genomic windows of size 1Mb, with a 500kb step size.

**S27**

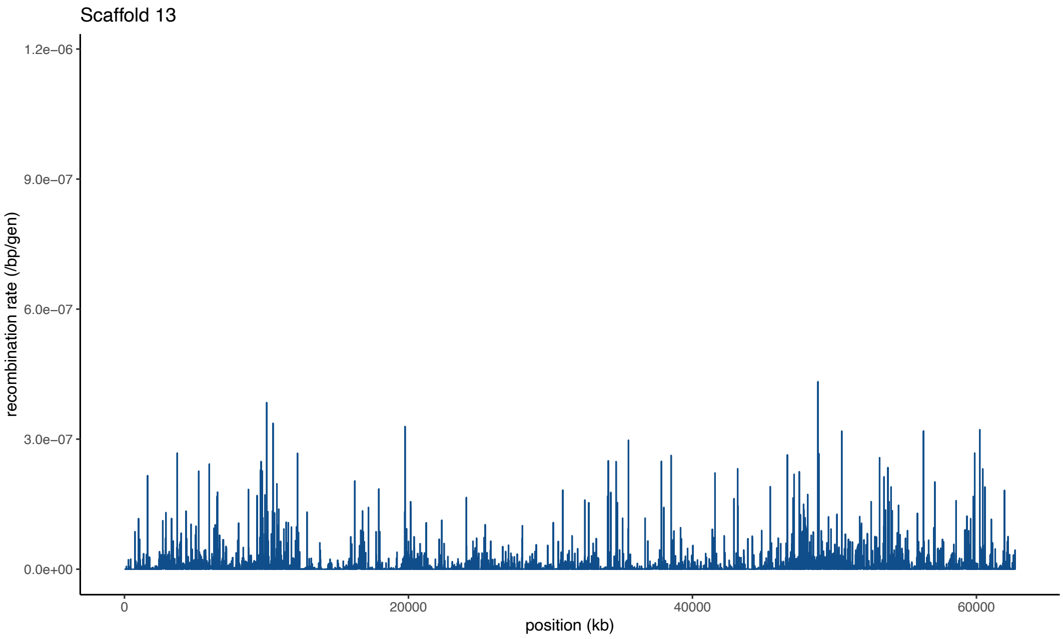


**Supplementary Figure S27:** Fine-scale per-base per-generation (/bp/gen) recombination rates along scaffold 13 for genomic windows of size 1Mb, with a 500kb step size.

**S28**

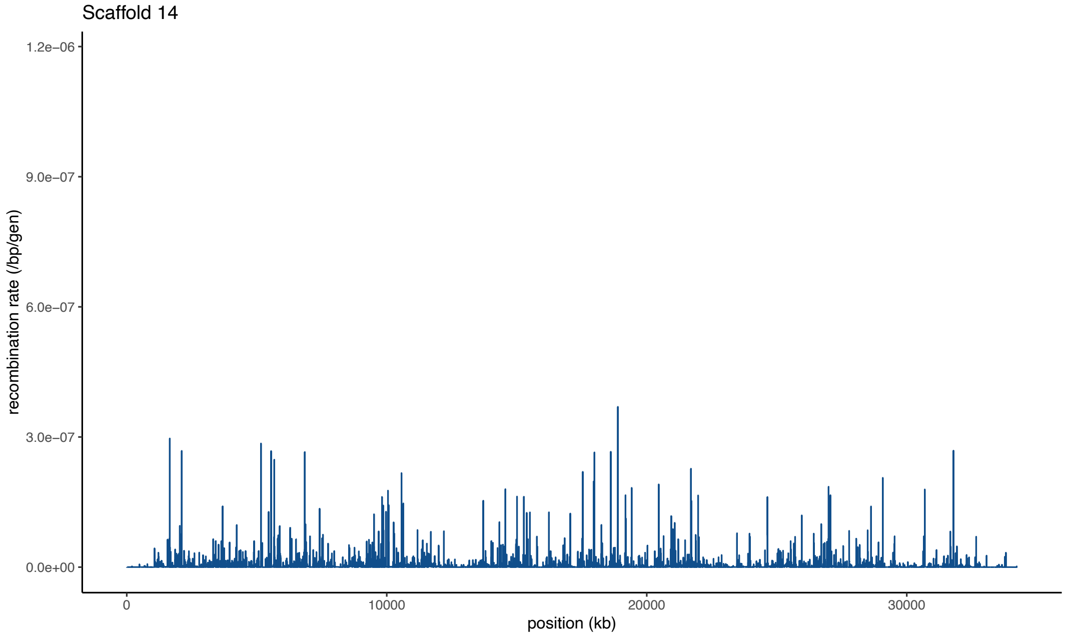


**Supplementary Figure S28:** Fine-scale per-base per-generation (/bp/gen) recombination rates along scaffold 14 for genomic windows of size 1Mb, with a 500kb step size.

**S29**

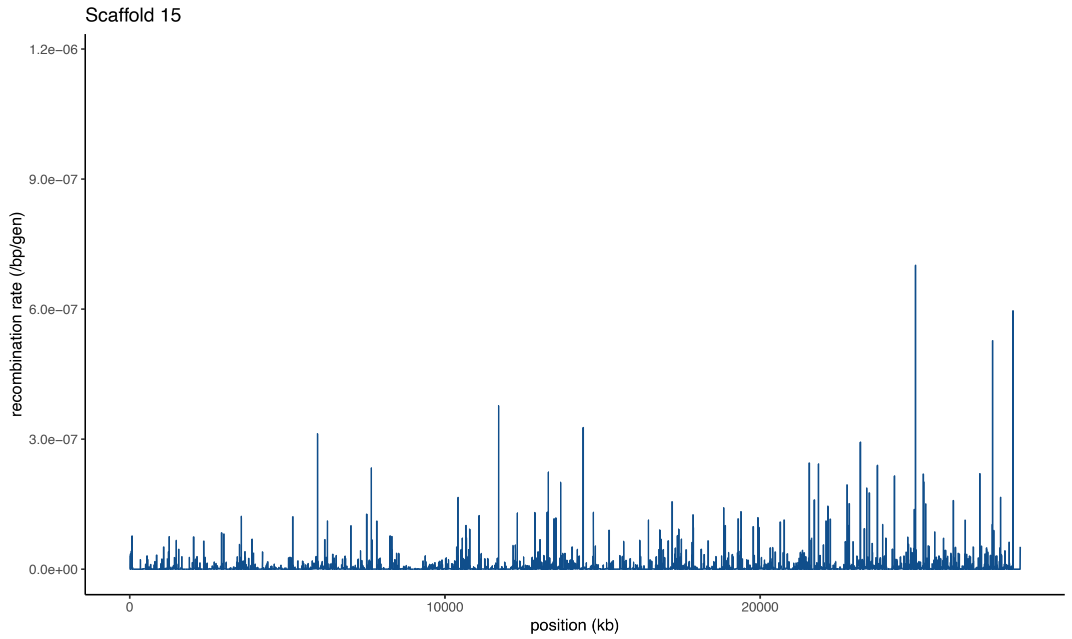


**Supplementary Figure S29:** Fine-scale per-base per-generation (/bp/gen) recombination rates along scaffold 15 for genomic windows of size 1Mb, with a 500kb step size.

## S30

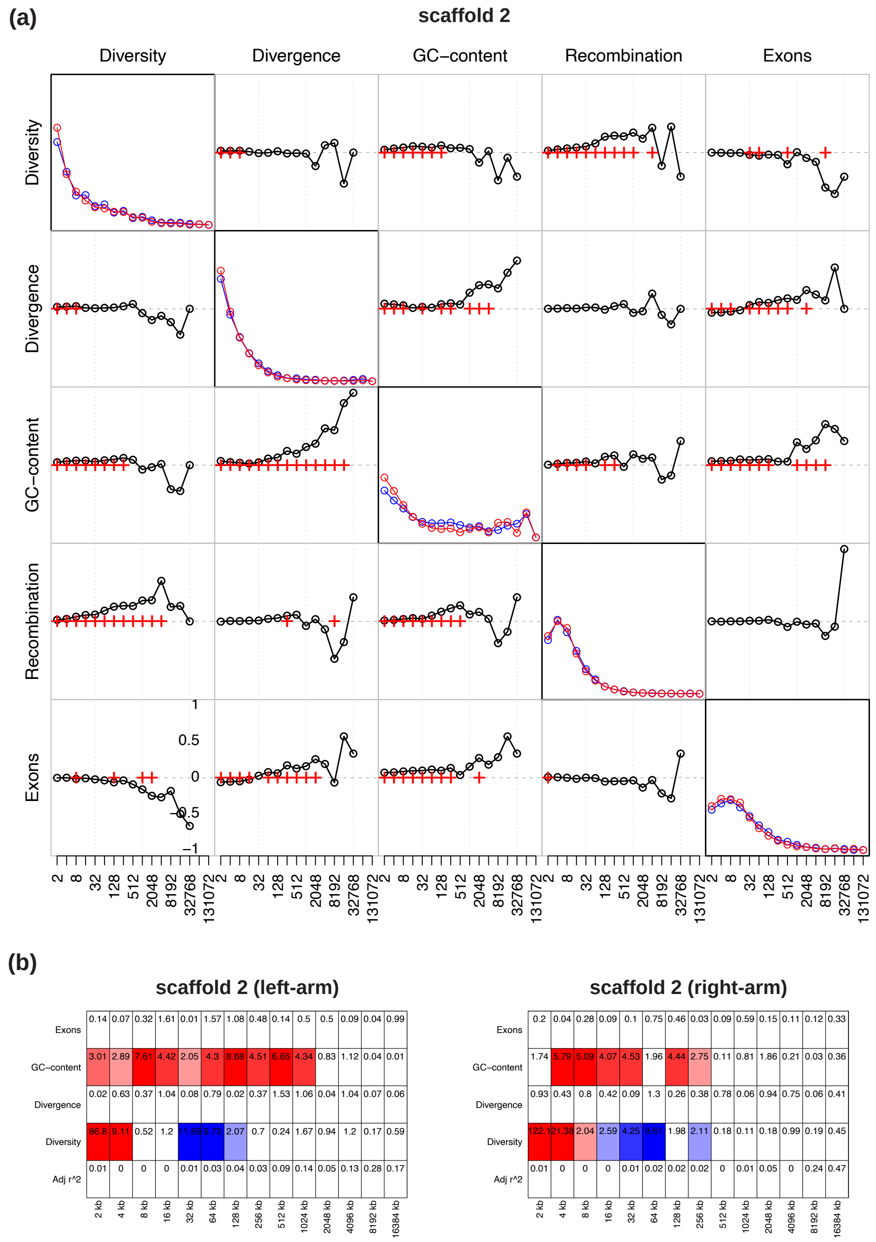


**Supplementary Figure S30:** (a) The detail coefficients of each genomic feature (diagonal plots) on the left and right arms of scaffold 2 (shown in dark and light yellow, respectively) as well as their pairwise correlations based on Kendall's rank correlation (off-diagonal plots with the bottom left showing the left-arm and the top right showing the right-arm) at a range of ($2^n$) scales. Correlations significant at the 1%-level under a two-tailed test are highlighted by crosses. (b) Linear model analysis of the detail coefficients. Red and blue coloring indicate significant positive and negative relationships under a two-sided *t*-test, with the color intensity being proportional to the significance level. Adjusted $r^2$ specifies the proportion of heterogeneity that can be explained by the linear model.

## S31

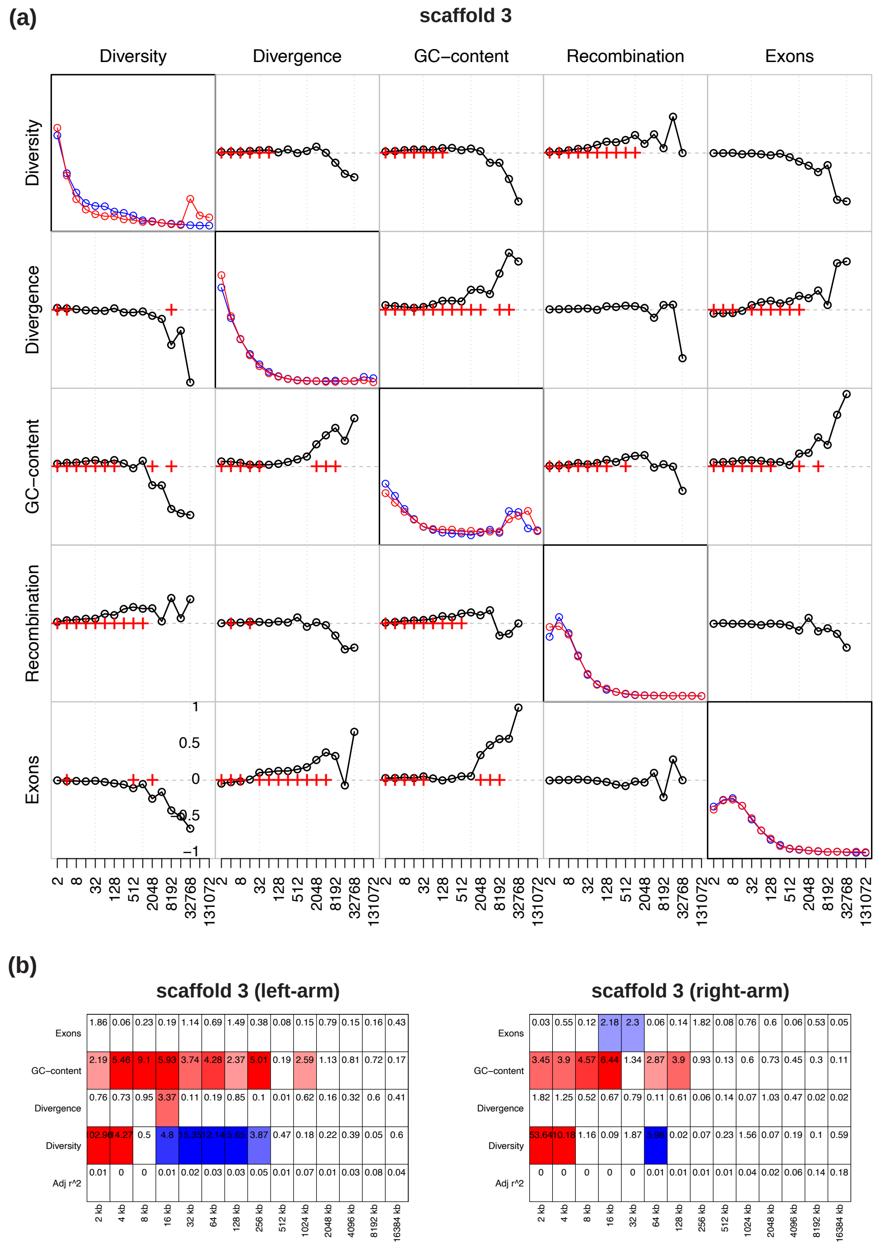


**Supplementary Figure S31:** (a) The detail coefficients of each genomic feature (diagonal plots) on the left and right arms of scaffold 3 (shown in dark and light yellow, respectively) as well as their pairwise correlations based on Kendall's rank correlation (off-diagonal plots with the bottom left showing the left-arm and the top right showing the right-arm) at a range of ($2^n$) scales. Correlations significant at the 1%-level under a two-tailed test are highlighted by crosses. (b) Linear model analysis of the detail coefficients. Red and blue coloring indicate significant positive and negative relationships under a two-sided *t*-test, with the color intensity being proportional to the significance level. Adjusted $r^2$ specifies the proportion of heterogeneity that can be explained by the linear model.

## S32

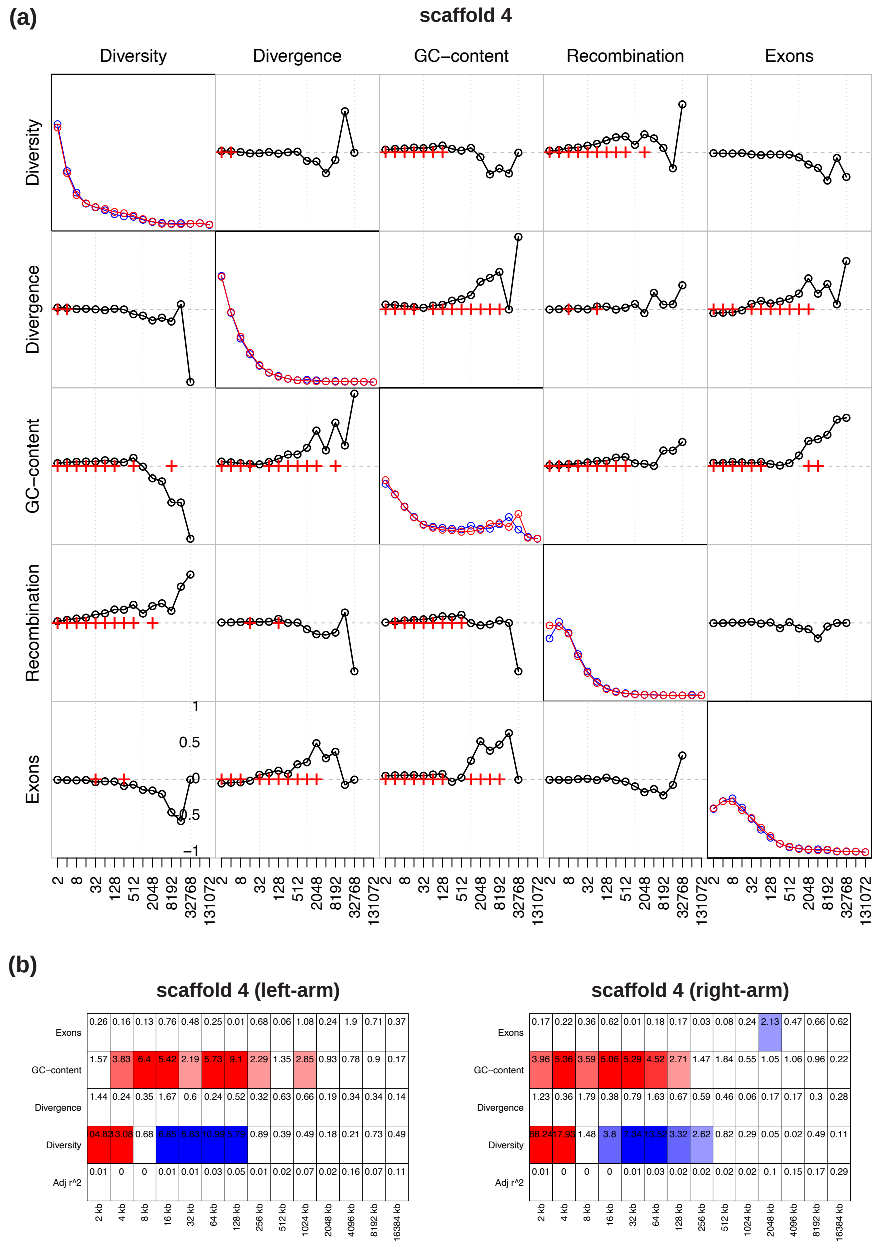


**Supplementary Figure S32:** (a) The detail coefficients of each genomic feature (diagonal plots) on the left and right arms of scaffold 4 (shown in dark and light yellow, respectively) as well as their pairwise correlations based on Kendall's rank correlation (off-diagonal plots with the bottom left showing the left-arm and the top right showing the right-arm) at a range of ($2^n$) scales. Correlations significant at the 1%-level under a two-tailed test are highlighted by crosses. (b) Linear model analysis of the detail coefficients. Red and blue coloring indicate significant positive and negative relationships under a two-sided *t*-test, with the color intensity being proportional to the significance level. Adjusted $r^2$ specifies the proportion of heterogeneity that can be explained by the linear model.

## S33

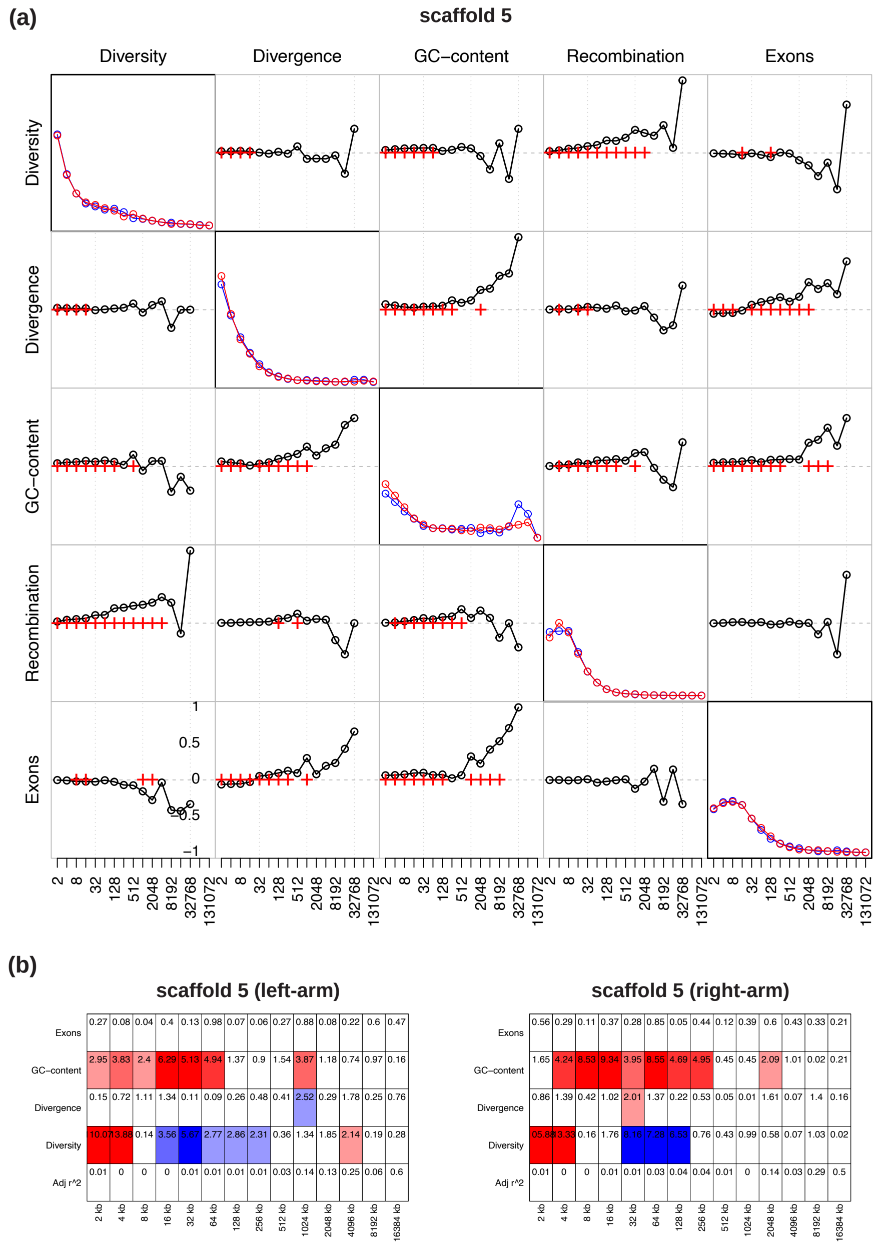


**Supplementary Figure S33:** (a) The detail coefficients of each genomic feature (diagonal plots) on the left and right arms of scaffold 5 (shown in dark and light yellow, respectively) as well as their pairwise correlations based on Kendall's rank correlation (off-diagonal plots with the bottom left showing the left-arm and the top right showing the right-arm) at a range of ($2^n$) scales. Correlations significant at the 1%-level under a two-tailed test are highlighted by crosses. (b) Linear model analysis of the detail coefficients. Red and blue coloring indicate significant positive and negative relationships under a two-sided *t*-test, with the color intensity being proportional to the significance level. Adjusted $r^2$ specifies the proportion of heterogeneity that can be explained by the linear model.

## S34

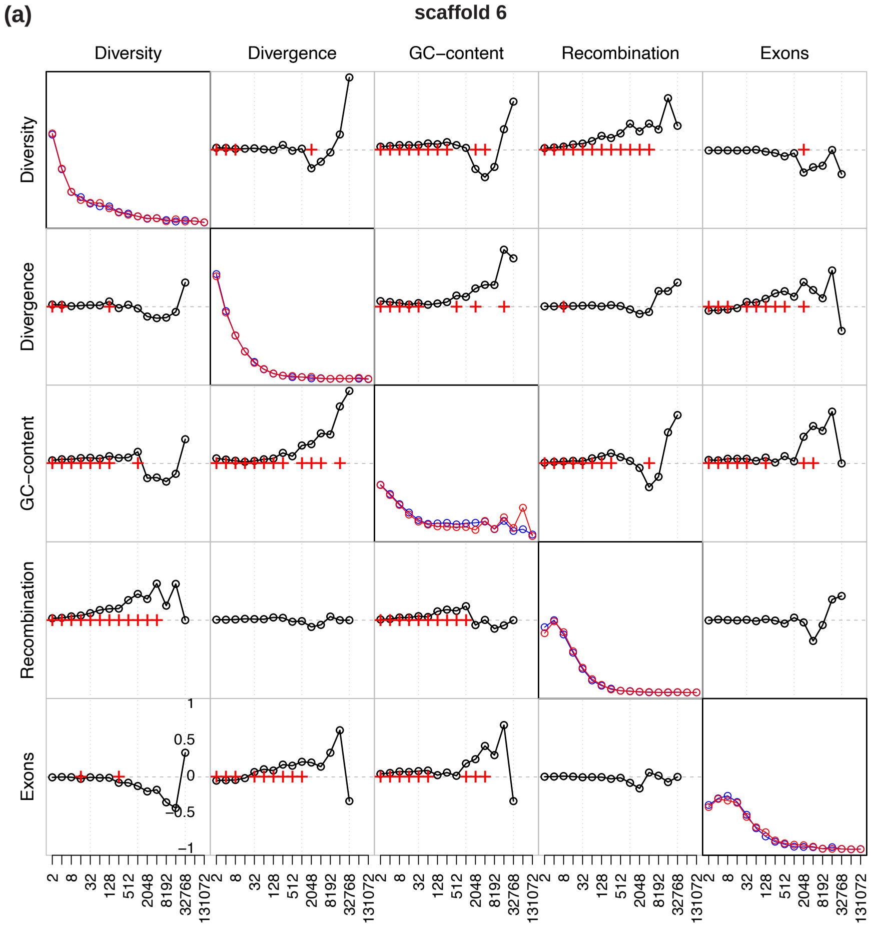


(b)

**scaffold 6 (left-arm)**

| | 2 kb | 4 kb | 8 kb | 16 kb | 32 kb | 64 kb | 128 kb | 256 kb | 512 kb | 1024 kb | 2048 kb | 4096 kb | 8192 kb | 16384 kb |
|---|---|---|---|---|---|---|---|---|---|---|---|---|---|---|
| Exons | 0.35 | 0.14 | 0.05 | 0.49 | 0.47 | 0.13 | 0.31 | 0.65 | 1.06 | 0.45 | 0.05 | 0.81 | 0.54 | 0.56 |
| GC−content | 3.06 | 7.62 | 9.41 | 7.31 | 7.18 | 1.98 | 5.26 | 7.49 | 2.91 | 2.15 | 0.54 | 0.36 | 0.28 | 0.06 |
| Divergence | 0.19 | 1.17 | 0.27 | 0.59 | 0.51 | 0.16 | 0.13 | 0.09 | 0.71 | 1.45 | 0.29 | 0.75 | 0.59 | 0.2 |
| Diversity | 10.15 | 11.2 | 2.3 | 4.38 | 10.18 | 6.12 | 5.72 | 1.59 | 0.15 | 1.7 | 2.67 | 1.85 | 0.2 | 1.26 |
| Adj r^2 | 0.01 | 0 | 0 | 0.01 | 0.02 | 0.01 | 0.03 | 0.06 | 0.04 | 0.13 | 0.11 | 0.1 | 0.03 | 0.39 |

**scaffold 6 (right-arm)**

| | 2 kb | 4 kb | 8 kb | 16 kb | 32 kb | 64 kb | 128 kb | 256 kb | 512 kb | 1024 kb | 2048 kb | 4096 kb | 8192 kb | 16384 kb |
|---|---|---|---|---|---|---|---|---|---|---|---|---|---|---|
| Exons | 0.36 | 2.34 | 0.26 | 0.05 | 0.06 | 0.22 | 0.06 | 0.37 | 0.5 | 0.15 | 0.1 | 0.1 | 0.3 | 0.05 |
| GC−content | 2.32 | 3.98 | 4.92 | 4.29 | 3.77 | 5.36 | 2.66 | 7.35 | 2.11 | 0.41 | 0.17 | 0.02 | 0.48 | 0.08 |
| Divergence | 1.14 | 0.94 | 0.45 | 0.13 | 0.53 | 0.16 | 0.02 | 0.52 | 1.83 | 0.2 | 0.08 | 0.31 | 0.53 | 0.19 |
| Diversity | 10.67 | 16.33 | 0.32 | 1.73 | 7.58 | 12.23 | 3.73 | 2.3 | 0.43 | 2.1 | 0.43 | 0.77 | 0.83 | 0.87 |
| Adj r^2 | 0.01 | 0 | 0 | 0 | 0.01 | 0.03 | 0.02 | 0.06 | 0.03 | 0.04 | 0.05 | 0.04 | 0.04 | 0.26 |

**Supplementary Figure S34:** (a) The detail coefficients of each genomic feature (diagonal plots) on the left and right arms of scaffold 6 (shown in dark and light yellow, respectively) as well as their pairwise correlations based on Kendall's rank correlation (off-diagonal plots with the bottom left showing the left-arm and the top right showing the right-arm) at a range of ($2^n$) scales. Correlations significant at the 1%-level under a two-tailed test are highlighted by crosses. (b) Linear model analysis of the detail coefficients. Red and blue coloring indicate significant positive and negative relationships under a two-sided *t*-test, with the color intensity being proportional to the significance level. Adjusted $r^2$ specifies the proportion of heterogeneity that can be explained by the linear model.

## S35

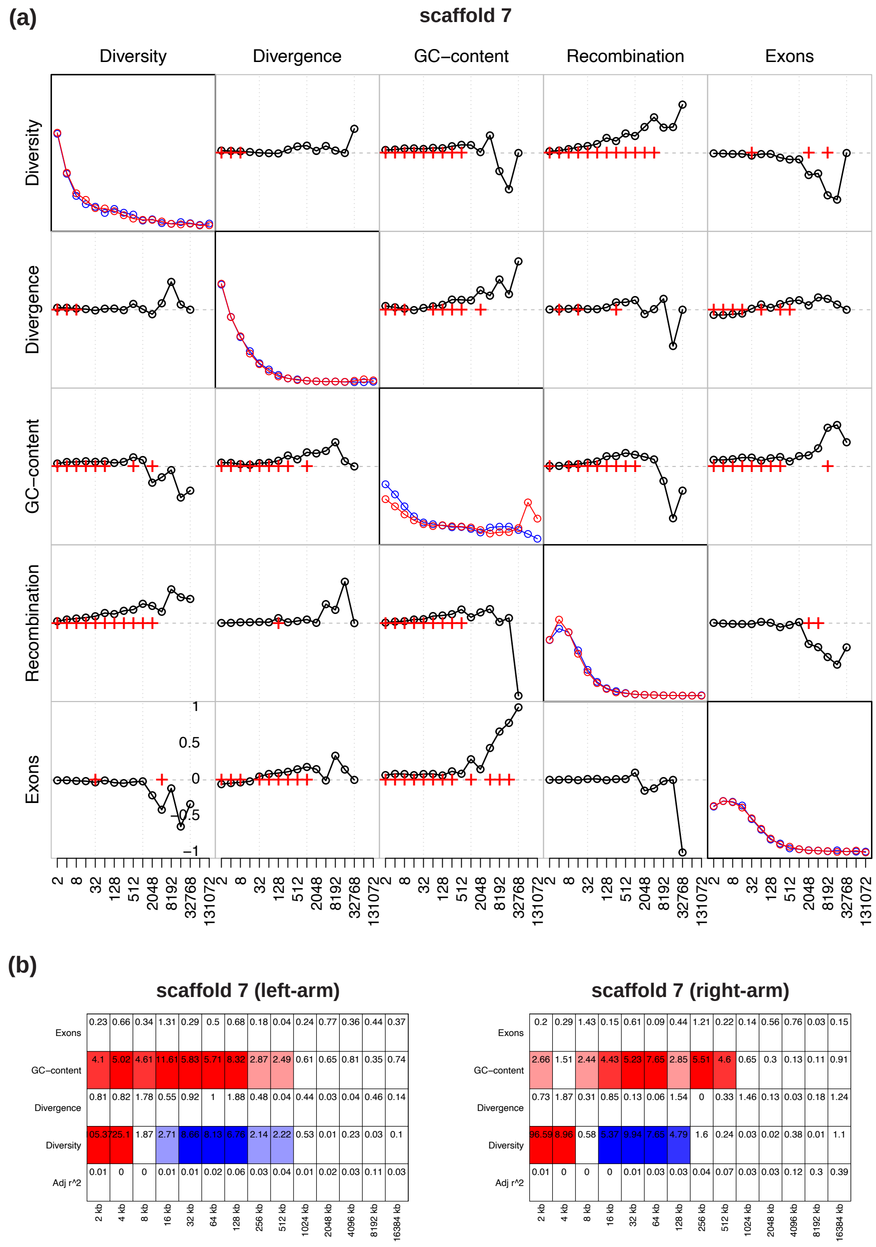


**Supplementary Figure S35:** (a) The detail coefficients of each genomic feature (diagonal plots) on the left and right arms of scaffold 7 (shown in dark and light yellow, respectively) as well as their pairwise correlations based on Kendall's rank correlation (off-diagonal plots with the bottom left showing the left-arm and the top right showing the right-arm) at a range of ($2^n$) scales. Correlations significant at the 1%-level under a two-tailed test are highlighted by crosses. (b) Linear model analysis of the detail coefficients. Red and blue coloring indicate significant positive and negative relationships under a two-sided *t*-test, with the color intensity being proportional to the significance level. Adjusted $r^2$ specifies the proportion of heterogeneity that can be explained by the linear model.

## S36

(a)

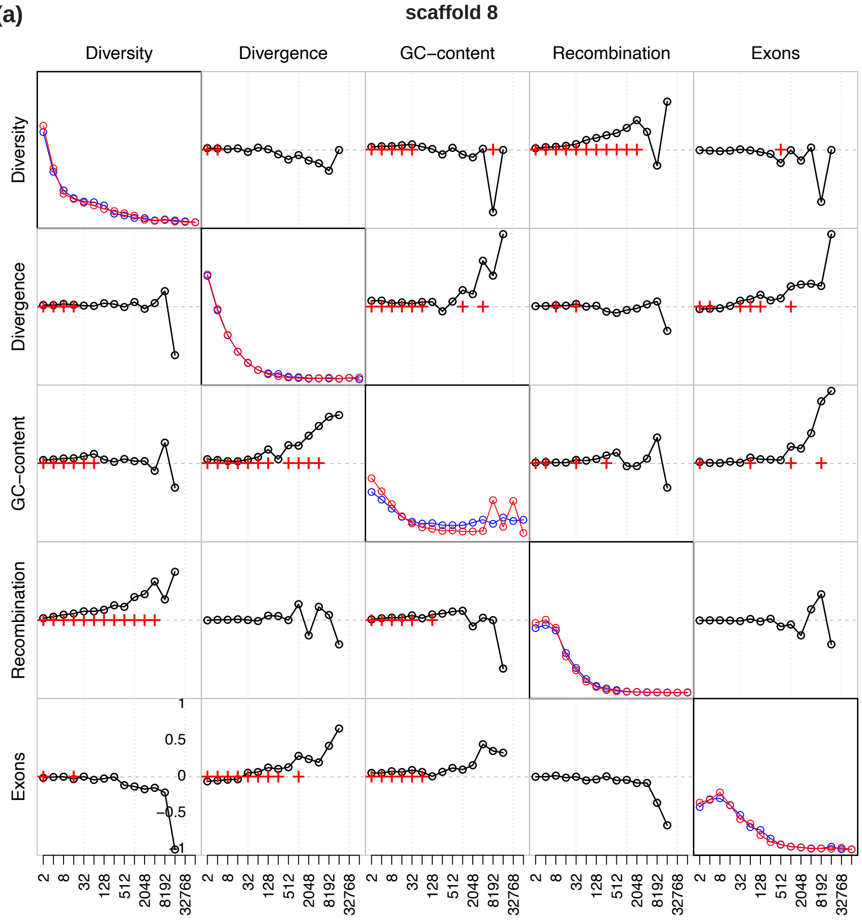


(b)

**scaffold 8 (left-arm)**

| | 2 kb | 4 kb | 8 kb | 16 kb | 32 kb | 64 kb | 128 kb | 256 kb | 512 kb | 1024 kb | 2048 kb | 4096 kb | 8192 kb |
|---|---|---|---|---|---|---|---|---|---|---|---|---|---|
| Exons | 0.27 | 1.78 | 0.4 | 0.6 | 0.45 | 2.29 | 0.4 | 0.49 | 0.55 | 0.98 | 0.46 | 0.48 | 0.55 |
| GC−content | 0.57 | 5.42 | 2.93 | 2.9 | 4.16 | 4.23 | 3.25 | 2.27 | 0.49 | 0.6 | 0.94 | 0.04 | 0.46 |
| Divergence | 0 | 0 | 0.42 | 0.38 | 0.64 | 0.47 | 0.19 | 0.63 | 0.12 | 1.97 | 0.23 | 0.31 | 0.01 |
| Diversity | 54.72 | 5.83 | 2.15 | 1.13 | 2.56 | 2.66 | 2.59 | 0.28 | 0.35 | 0.15 | 1.38 | 0.88 | 0.09 |
| Adj r^2 | 0.01 | 0 | 0 | 0 | 0.01 | 0.02 | 0.03 | 0.02 | 0.01 | 0.12 | 0.21 | 0.02 | 0.1 |

**scaffold 8 (right-arm)**

| | 2 kb | 4 kb | 8 kb | 16 kb | 32 kb | 64 kb | 128 kb | 256 kb | 512 kb | 1024 kb | 2048 kb | 4096 kb | 8192 kb |
|---|---|---|---|---|---|---|---|---|---|---|---|---|---|
| Exons | 0.47 | 0.08 | 0.41 | 0.65 | 0.37 | 0.04 | 0.36 | 0.91 | 0.75 | 0.22 | 0.53 | 0.01 | 0.01 |
| GC−content | 3.31 | 4.13 | 1.58 | 0.06 | 2.45 | 1.06 | 0.71 | 1.1 | 0.69 | 0.19 | 0.45 | 0.09 | 0.08 |
| Divergence | 0.13 | 0.83 | 0.38 | 0.61 | 0.87 | 0.12 | 0.03 | 0.09 | 0.2 | 0.02 | 0.06 | 0.45 | 0.11 |
| Diversity | 64.48 | 12.66 | 0.32 | 0.8 | 0.97 | 0.36 | 0.22 | 0.24 | 0.25 | 0.39 | 0.89 | 0.03 | 0.28 |
| Adj r^2 | 0.01 | 0 | 0 | 0 | 0 | 0 | 0 | 0.01 | 0 | 0.04 | 0.06 | 0.18 | 0.59 |

**Supplementary Figure S36:** (a) The detail coefficients of each genomic feature (diagonal plots) on the left and right arms of scaffold 8 (shown in dark and light yellow, respectively) as well as their pairwise correlations based on Kendall's rank correlation (off-diagonal plots with the bottom left showing the left-arm and the top right showing the right-arm) at a range of ($2^n$) scales. Correlations significant at the 1%-level under a two-tailed test are highlighted by crosses. (b) Linear model analysis of the detail coefficients. Red and blue coloring indicate significant positive and negative relationships under a two-sided *t*-test, with the color intensity being proportional to the significance level. Adjusted $r^2$ specifies the proportion of heterogeneity that can be explained by the linear model.

## S37

(a)

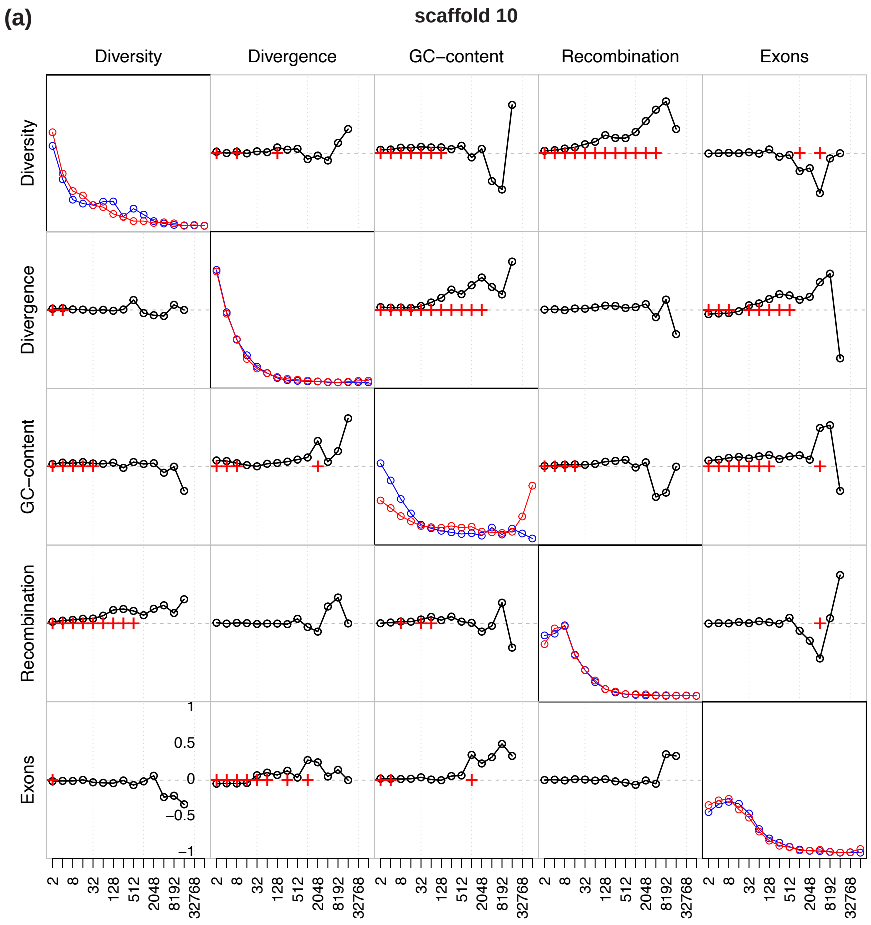


(b)

**scaffold 10 (left-arm)**

| | 2 kb | 4 kb | 8 kb | 16 kb | 32 kb | 64 kb | 128 kb | 256 kb | 512 kb | 1024 kb | 2048 kb | 4096 kb | 8192 kb |
|---|---|---|---|---|---|---|---|---|---|---|---|---|---|
| Exons | 0.89 | 1.15 | 0.05 | 0.12 | 0.03 | 0.01 | 0.86 | 0.09 | 0.26 | 0.28 | 0.32 | 0.23 | 0.46 |
| GC−content | 0.08 | 5.25 | 2.79 | 1.52 | 2.1 | 3.35 | 2.01 | 5.33 | 0.88 | 0.28 | 0.24 | 0.05 | 0.51 |
| Divergence | 0.86 | 0.33 | 0.99 | 0.05 | 0.53 | 0.14 | 0.4 | 0.21 | 0.76 | 0.17 | 0.64 | 0.21 | 0.5 |
| Diversity | 65.75 | 6.39 | 0.7 | 1.3 | 4.87 | 3.57 | 0.27 | 0.11 | 0.64 | 0.59 | 0.11 | 0.16 | 0.04 |
| Adj r^2 | 0.01 | 0 | 0 | 0 | 0.01 | 0.02 | 0.01 | 0.07 | 0.01 | 0.01 | 0.06 | 0.27 | 0.17 |

**scaffold 10 (right-arm)**

| | 2 kb | 4 kb | 8 kb | 16 kb | 32 kb | 64 kb | 128 kb | 256 kb | 512 kb | 1024 kb | 2048 kb | 4096 kb | 8192 kb |
|---|---|---|---|---|---|---|---|---|---|---|---|---|---|
| Exons | 0.06 | 0.76 | 0.64 | 0.29 | 0.41 | 0.03 | 0.42 | 0.66 | 0.41 | 0.09 | 0.14 | 0.21 | 0.11 |
| GC−content | 3.84 | 1.19 | 2.24 | 1.27 | 0.5 | 0.84 | 6.06 | 1.25 | 1.63 | 1.26 | 0.03 | 0.48 | 0.23 |
| Divergence | 0.67 | 0.3 | 0.49 | 0.62 | 0.08 | 0.68 | 0.49 | 0.19 | 0.27 | 0.46 | 0.02 | 0.2 | 0.31 |
| Diversity | 46.96 | 9.24 | 0.05 | 0.59 | 1.23 | 2.23 | 1.18 | 2.09 | 1.41 | 1.47 | 0.98 | 1.26 | 0.37 |
| Adj r^2 | 0.01 | 0 | 0 | 0 | 0 | 0.01 | 0.04 | 0.03 | 0.09 | 0.08 | 0.01 | 0.07 | 0.22 |

**Supplementary Figure S37:** (a) The detail coefficients of each genomic feature (diagonal plots) on the left and right arms of scaffold 10 (shown in dark and light yellow, respectively) as well as their pairwise correlations based on Kendall's rank correlation (off-diagonal plots with the bottom left showing the left-arm and the top right showing the right-arm) at a range of ($2^n$) scales. Correlations significant at the 1%-level under a two-tailed test are highlighted by crosses. (b) Linear model analysis of the detail coefficients. Red and blue coloring indicate significant positive and negative relationships under a two-sided *t*-test, with the color intensity being proportional to the significance level. Adjusted $r^2$ specifies the proportion of heterogeneity that can be explained by the linear model.

## S38

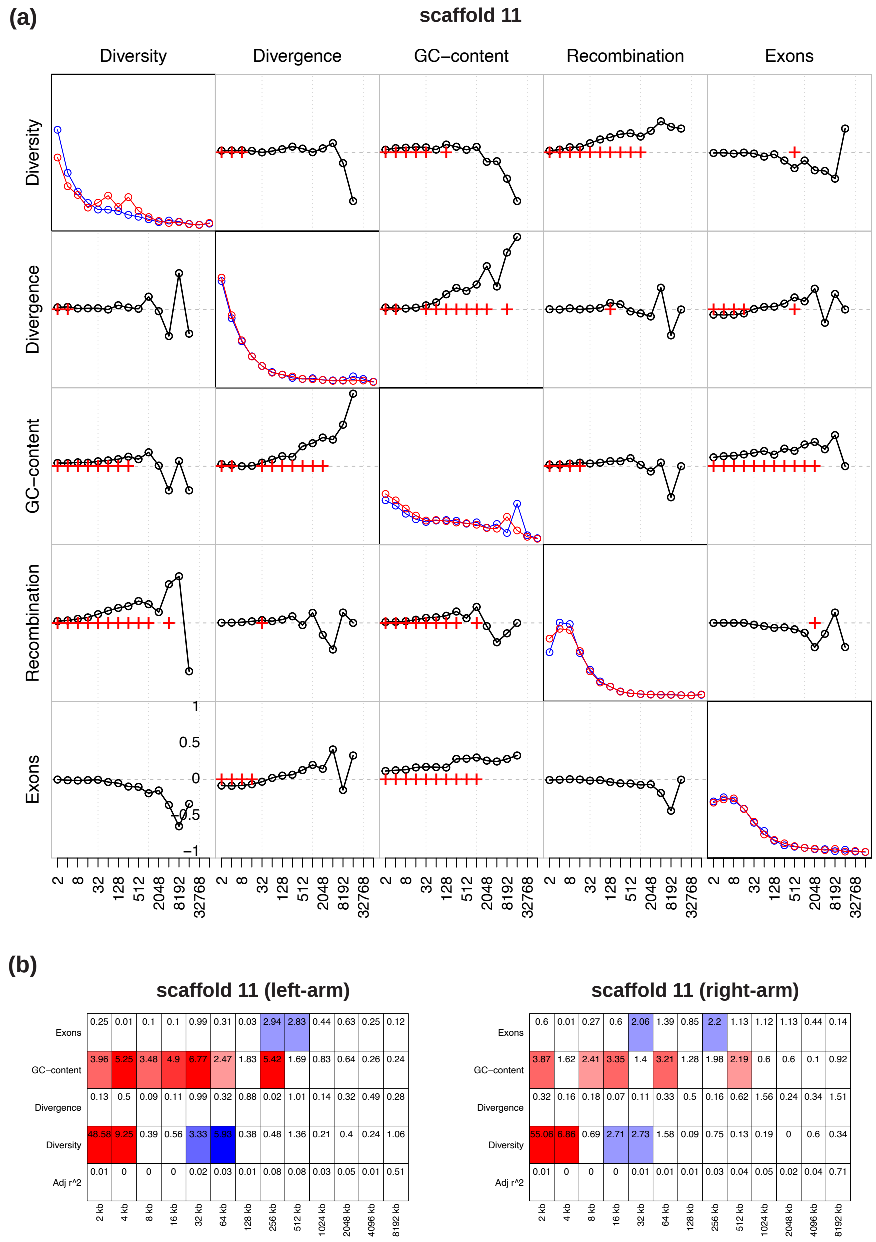


**Supplementary Figure S38:** (a) The detail coefficients of each genomic feature (diagonal plots) on the left and right arms of scaffold 11 (shown in dark and light yellow, respectively) as well as their pairwise correlations based on Kendall's rank correlation (off-diagonal plots with the bottom left showing the left-arm and the top right showing the right-arm) at a range of ($2^n$) scales. Correlations significant at the 1%-level under a two-tailed test are highlighted by crosses. (b) Linear model analysis of the detail coefficients. Red and blue coloring indicate significant positive and negative relationships under a two-sided *t*-test, with the color intensity being proportional to the significance level. Adjusted $r^2$ specifies the proportion of heterogeneity that can be explained by the linear model.

## S39

(a)

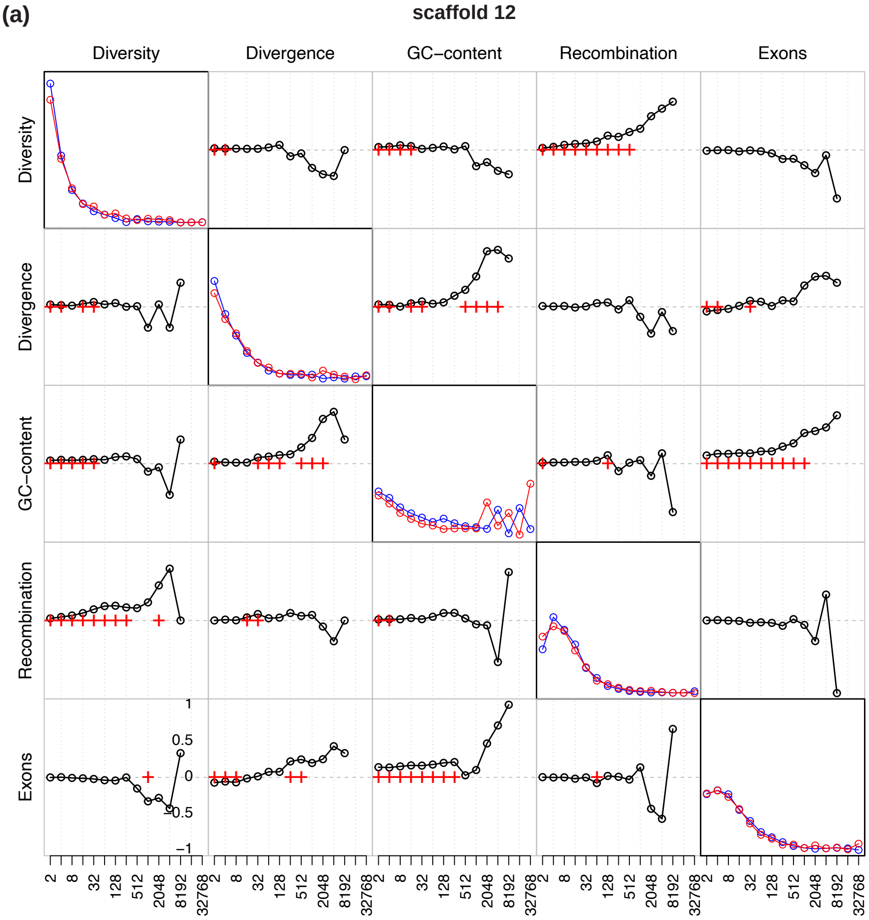


(b)

**scaffold 12 (left-arm)**

| | 2 kb | 4 kb | 8 kb | 16 kb | 32 kb | 64 kb | 128 kb | 256 kb | 512 kb | 1024 kb | 2048 kb | 4096 kb |
|---|---|---|---|---|---|---|---|---|---|---|---|---|
| Exons | 0.67 | 0.04 | 0.39 | 1.16 | 0.17 | 1.07 | 0.68 | 0.37 | 0.79 | 0.05 | 0 | 0.26 |
| GC−content | 0.73 | 2.09 | 0.36 | 0.35 | 1.52 | 1.31 | 2.45 | 1.35 | 0.13 | 0.29 | 0.18 | 0.46 |
| Divergence | 0.14 | 0.58 | 0.52 | 0.2 | 0.51 | 0.14 | 0.53 | 0.19 | 0.2 | 0.23 | 0.23 | 0.66 |
| Diversity | 26.26 | 2.92 | 0.09 | 2.11 | 0.71 | 0.62 | 0.28 | 1.12 | 0.58 | 1.03 | 0.84 | 0.8 |
| Adj r^2 | 0.01 | 0 | 0 | 0 | 0 | 0.01 | 0.03 | 0.04 | 0.01 | 0.02 | 0.09 | 0.2 |

**scaffold 12 (right-arm)**

| | 2 kb | 4 kb | 8 kb | 16 kb | 32 kb | 64 kb | 128 kb | 256 kb | 512 kb | 1024 kb | 2048 kb | 4096 kb |
|---|---|---|---|---|---|---|---|---|---|---|---|---|
| Exons | 0.56 | 0.33 | 0 | 0.74 | 0.09 | 0.16 | 0.1 | 0.52 | 0.72 | 0.16 | 0.28 | 0 |
| GC−content | 0.69 | 0.73 | 0.83 | 1.7 | 1.25 | 0.87 | 1.01 | 0.02 | 0.32 | 0.49 | 0.65 | 0.43 |
| Divergence | 0.09 | 0.36 | 0.18 | 0.02 | 0.17 | 0.38 | 0.2 | 0.45 | 0.21 | 0.75 | 1.16 | 0.01 |
| Diversity | 31.45 | 5.14 | 0.08 | 0.46 | 1.75 | 0.96 | 0.19 | 0.39 | 0.84 | 0 | 0.2 | 1.46 |
| Adj r^2 | 0.01 | 0 | 0 | 0 | 0.01 | 0 | 0 | 0.01 | 0.03 | 0.06 | 0.07 | 0.66 |

**Supplementary Figure S39:** (a) The detail coefficients of each genomic feature (diagonal plots) on the left and right arms of scaffold 12 (shown in dark and light yellow, respectively) as well as their pairwise correlations based on Kendall's rank correlation (off-diagonal plots with the bottom left showing the left-arm and the top right showing the right-arm) at a range of ($2^n$) scales. Correlations significant at the 1%-level under a two-tailed test are highlighted by crosses. (b) Linear model analysis of the detail coefficients. Red and blue coloring indicate significant positive and negative relationships under a two-sided *t*-test, with the color intensity being proportional to the significance level. Adjusted $r^2$ specifies the proportion of heterogeneity that can be explained by the linear model.

## S40

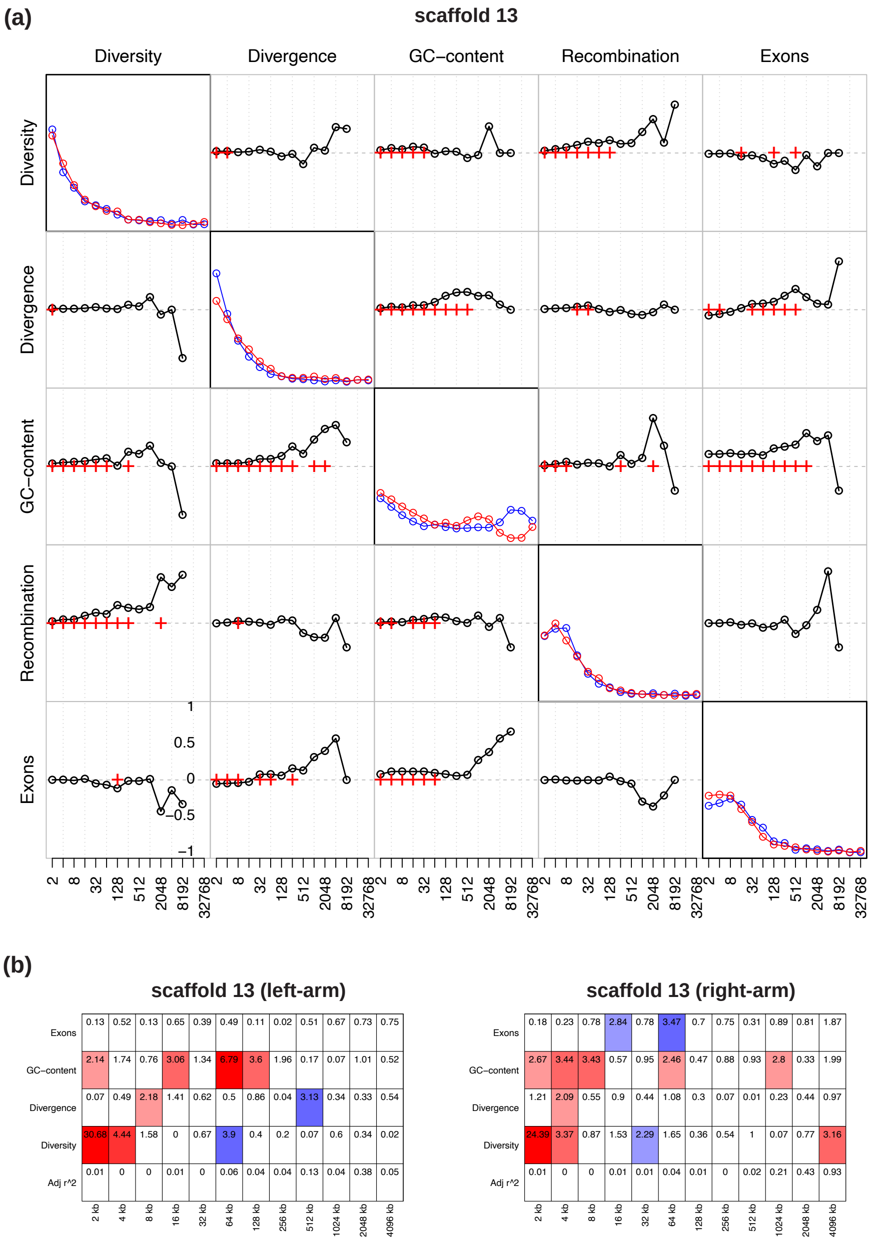


**Supplementary Figure S40:** (a) The detail coefficients of each genomic feature (diagonal plots) on the left and right arms of scaffold 13 (shown in dark and light yellow, respectively) as well as their pairwise correlations based on Kendall's rank correlation (off-diagonal plots with the bottom left showing the left-arm and the top right showing the right-arm) at a range of ($2^n$) scales. Correlations significant at the 1%-level under a two-tailed test are highlighted by crosses. (b) Linear model analysis of the detail coefficients. Red and blue coloring indicate significant positive and negative relationships under a two-sided *t*-test, with the color intensity being proportional to the significance level. Adjusted $r^2$ specifies the proportion of heterogeneity that can be explained by the linear model.

## S41

(a)

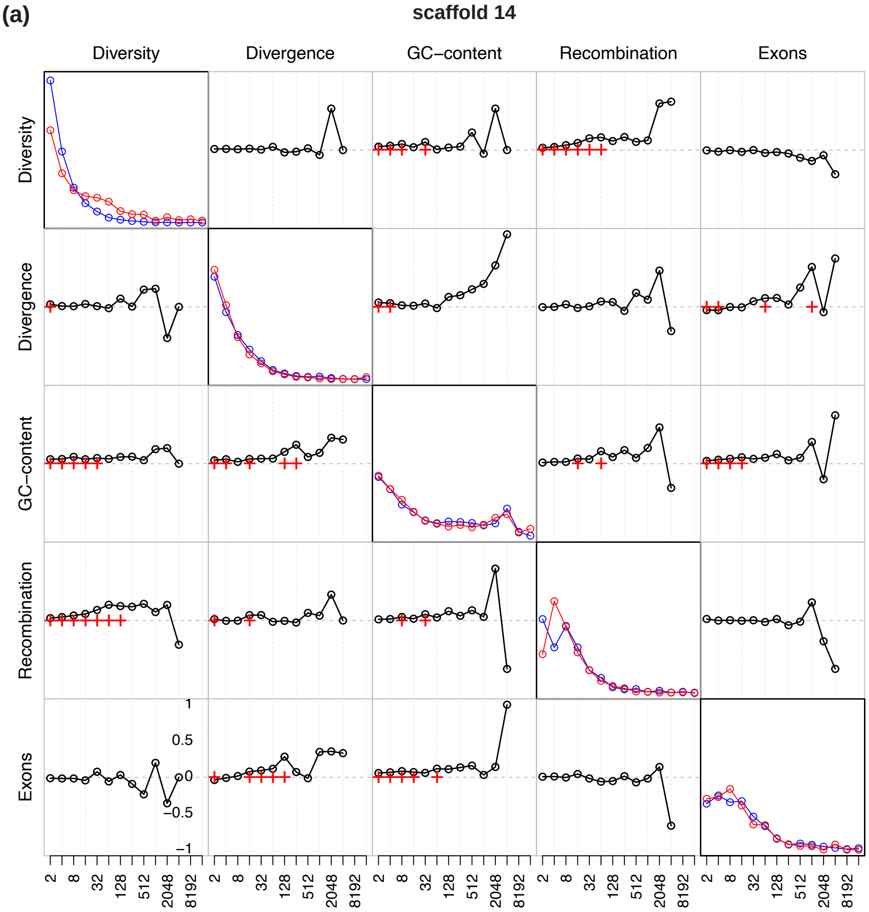


(b)

**scaffold 14 (left-arm)**

| | 2 kb | 4 kb | 8 kb | 16 kb | 32 kb | 64 kb | 128 kb | 256 kb | 512 kb | 1024 kb | 2048 kb |
|---|---|---|---|---|---|---|---|---|---|---|---|
| Exons | 0.31 | 0.27 | 0.04 | 0.41 | 2.38 | 0.82 | 0.13 | 0.02 | 0.65 | 0.17 | 0.65 |
| GC−content | 0.24 | 1.58 | 4.18 | 1.19 | 2.5 | 0.59 | 0.3 | 1.5 | 0.08 | 0.14 | 0.46 |
| Divergence | 1.1 | 0.03 | 1.18 | 0.93 | 0.81 | 0.63 | 0.29 | 2.88 | 0.24 | 0.09 | 0.2 |
| Diversity | 13.7 | 0.83 | 0.44 | 1.17 | 0.82 | 0.65 | 0.58 | 0.13 | 0.71 | 0.36 | 1.15 |
| Adj r^2 | 0.01 | 0 | 0.01 | 0.01 | 0.03 | 0.01 | 0.01 | 0.12 | 0.09 | 0.23 | 0.6 |

**scaffold 14 (right-arm)**

| | 2 kb | 4 kb | 8 kb | 16 kb | 32 kb | 64 kb | 128 kb | 256 kb | 512 kb | 1024 kb | 2048 kb |
|---|---|---|---|---|---|---|---|---|---|---|---|
| Exons | 0.02 | 0.25 | 0.1 | 0.09 | 0.51 | 0.71 | 0.09 | 0.65 | 0.54 | 0.22 | 0.67 |
| GC−content | 2.32 | 2.54 | 1.91 | 2.1 | 1.23 | 0.83 | 1.3 | 0.65 | 1.9 | 1.21 | 0.19 |
| Divergence | 0.59 | 0.39 | 0.08 | 0.16 | 0.02 | 1.48 | 0.47 | 0.12 | 0.23 | 0.41 | 0.3 |
| Diversity | 15.19 | 1.65 | 0.02 | 0.79 | 0.74 | 1.71 | 0.9 | 0.12 | 0.7 | 0.55 | 0.07 |
| Adj r^2 | 0.01 | 0 | 0 | 0 | 0 | 0.03 | 0.04 | 0.01 | 0.16 | 0.2 | 0.13 |

**Supplementary Figure S41:** (a) The detail coefficients of each genomic feature (diagonal plots) on the left and right arms of scaffold 14 (shown in dark and light yellow, respectively) as well as their pairwise correlations based on Kendall's rank correlation (off-diagonal plots with the bottom left showing the left-arm and the top right showing the right-arm) at a range of ($2^n$) scales. Correlations significant at the 1%-level under a two-tailed test are highlighted by crosses. (b) Linear model analysis of the detail coefficients. Red and blue coloring indicate significant positive and negative relationships under a two-sided *t*-test, with the color intensity being proportional to the significance level. Adjusted $r^2$ specifies the proportion of heterogeneity that can be explained by the linear model.

## S42

(a)

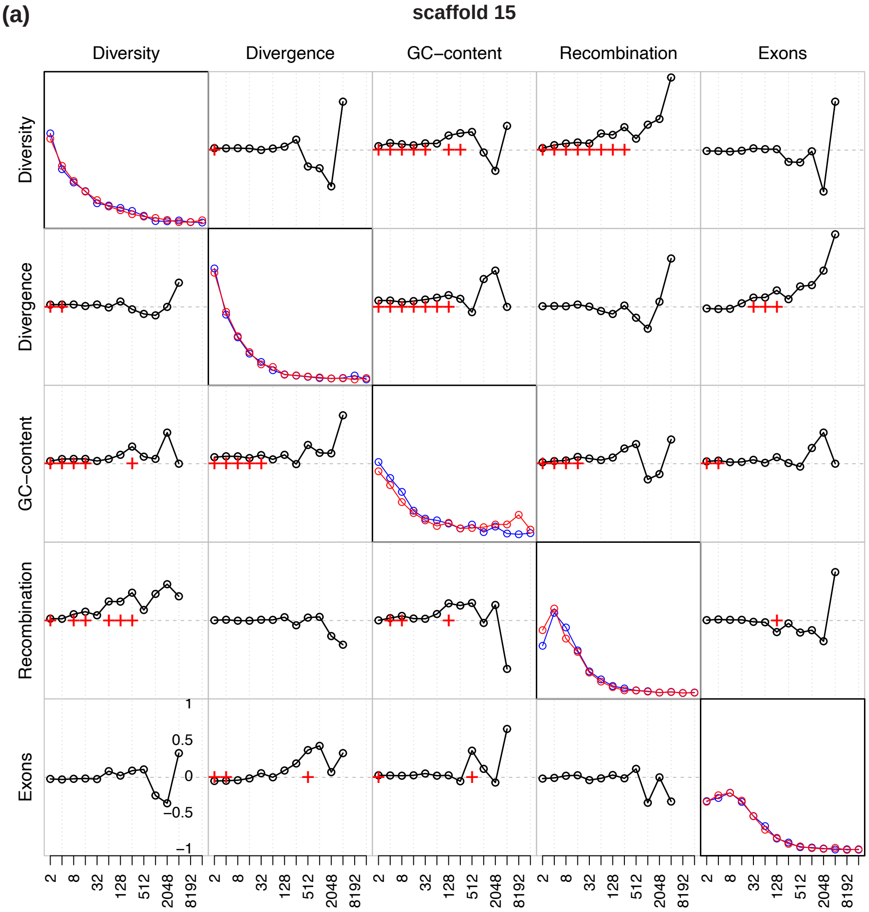


(b)

**scaffold 15 (left-arm)**

| | 2 kb | 4 kb | 8 kb | 16 kb | 32 kb | 64 kb | 128 kb | 256 kb | 512 kb | 1024 kb | 2048 kb |
|---|---|---|---|---|---|---|---|---|---|---|---|
| Exons | 1.33 | 0.69 | 1.13 | 0.03 | 0.45 | 0.52 | 0.22 | 0.33 | 0.02 | 0.02 | 0.2 |
| GC−content | 0.49 | 2.8 | 2.57 | 0.21 | 0.85 | 1.45 | 0.91 | 0.25 | 2.22 | 0.31 | 0.25 |
| Divergence | 0.35 | 0.31 | 0.45 | 0.78 | 0.01 | 0.75 | 0.68 | 1.56 | 0.84 | 0.15 | 0.02 |
| Diversity | 11.31 | 0.45 | 0.7 | 1.36 | 4.68 | 0.76 | 0.93 | 0.27 | 1.06 | 0 | 0.27 |
| Adj r^2 | 0.01 | 0 | 0 | 0 | 0.03 | 0.02 | 0.02 | 0.03 | 0.31 | 0.27 | 0.14 |

**scaffold 15 (right-arm)**

| | 2 kb | 4 kb | 8 kb | 16 kb | 32 kb | 64 kb | 128 kb | 256 kb | 512 kb | 1024 kb | 2048 kb |
|---|---|---|---|---|---|---|---|---|---|---|---|
| Exons | 0.55 | 0.62 | 0.13 | 0.4 | 0.41 | 0.11 | 0.61 | 0.3 | 0.41 | 0.72 | 0.07 |
| GC−content | 1.58 | 2.24 | 4.47 | 4.34 | 3.59 | 1.53 | 1.5 | 2.23 | 1.42 | 0.22 | 0.26 |
| Divergence | 0.15 | 0 | 0.8 | 1.33 | 0.01 | 0.09 | 0.8 | 0.39 | 0.16 | 0.95 | 0.26 |
| Diversity | 14.39 | 1.43 | 0.16 | 2.59 | 2.79 | 0.54 | 0.02 | 0.26 | 1.18 | 1.8 | 1.12 |
| Adj r^2 | 0.01 | 0 | 0.01 | 0.02 | 0.03 | 0.01 | 0.03 | 0.09 | 0.3 | 0.31 | 0.41 |

**Supplementary Figure S42:** (a) The detail coefficients of each genomic feature (diagonal plots) on the left and right arms of scaffold 15 (shown in dark and light yellow, respectively) as well as their pairwise correlations based on Kendall's rank correlation (off-diagonal plots with the bottom left showing the left-arm and the top right showing the right-arm) at a range of ($2^n$) scales. Correlations significant at the 1%-level under a two-tailed test are highlighted by crosses. (b) Linear model analysis of the detail coefficients. Red and blue coloring indicate significant positive and negative relationships under a two-sided *t*-test, with the color intensity being proportional to the significance level. Adjusted $r^2$ specifies the proportion of heterogeneity that can be explained by the linear model.
